# Intranasal Vaccination with a Lentiviral Vector Strongly Protects against SARS-CoV-2 in Mouse and Golden Hamster Preclinical Models

**DOI:** 10.1101/2020.07.21.214049

**Authors:** Min-Wen Ku, Maryline Bourgine, Pierre Authié, Jodie Lopez, Kirill Nemirov, Fanny Moncoq, Amandine Noirat, Benjamin Vesin, Fabien Nevo, Catherine Blanc, Philippe Souque, Houda Tabbal, Emeline Simon, Marine Le Dudal, Françoise Guinet, Laurence Fiette, Hugo Mouquet, François Anna, Annette Martin, Nicolas Escriou, Laleh Majlessi, Pierre Charneau

## Abstract

To develop a vaccine candidate against COVID-19, we generated a Lentiviral Vector (LV), eliciting neutralizing antibodies against the Spike glycoprotein of SARS-CoV-2. Systemic vaccination by this vector in mice, in which the expression of the SARS-CoV-2 receptor hACE2 has been induced by transduction of respiratory tract cells by an adenoviral vector, conferred only partial protection, despite an intense serum neutralizing activity. However, targeting the immune response to the respiratory tract through an intranasal boost with this LV resulted in > 3 log10 decrease in the lung viral loads and avoided local inflammation. Moreover, both integrative and non-integrative LV platforms displayed a strong vaccine efficacy and inhibited lung deleterious injury in golden hamsters, which are naturally permissive to SARS-CoV-2 replication and restitute the human COVID-19 physiopathology. Our results provide evidence of marked prophylactic effects of the LV-based vaccination against SARS-CoV-2 and designate the intranasal immunization as a powerful approach against COVID-19.

**Highlights:** A lentiviral vector encoding for Spike predicts a promising COVID-19 vaccine

Targeting the immune response to the upper respiratory tract is key to protection

Intranasal vaccination induces protective mucosal immunity against SARS-CoV-2

Lung anti-Spike IgA responses correlate with protection and reduced inflammation

## Introduction

The new Severe Acute Respiratory Syndrome beta-coronavirus 2 (SARS-CoV-2) that emerged in late 2019 in Wuhan, China, is extraordinarily contagious and fast-spreading across the world (Guo et al., 2020). Compared to the previously emerged SARS or Middle East Respiratory Syndrome (MERS) coronaviruses, SARS-CoV-2 causes unprecedented threat on global health and tremendous socio-economic consequences. Therefore, the development of effective prophylactic vaccines against SARS-CoV-2 is an absolute imperative to contain the spread of the epidemic and to prevent the development of CoronaVirus Disease 2019 (COVID-19)-associated symptoms such as deleterious inflammation and progressive respiratory failure (Amanat and Krammer, 2020).

Coronaviruses are enveloped, non-segmented positive-stranded RNA viruses, characterized by their envelope-anchored Spike (S) glycoprotein (Walls et al., 2020). The SARS-CoV-2 S (S_CoV-2_) is a (180 kDa)_3_ homotrimeric class I viral fusion protein, that engages the carboxypeptidase Angiotensin-Converting Enzyme 2 (ACE2) expressed on host cells. The monomer of S_CoV-2_ protein possesses an ectodomain, a transmembrane anchor domain, and a short internal tail. S_CoV-2_ is activated by a two-step sequential proteolytic cleavage to initiate fusion with the host cell membrane. Subsequent to S_CoV-2_-ACE2 interaction, through a conformational reorganization, the extracellular domain of S_CoV-2_ is first cleaved at the highly specific furin 682^RRAR^685 site (Guo et al., 2020; Walls et al., 2020), a key factor determining the pathological features of the virus, linked to the furin ubiquitous expression (Wang et al., 2020). The resulting subunits are: (i) S1, which harbors the ACE2 Receptor Binding Domain (RBD), with the atomic contacts restricted to the ACE2 protease domain, and (ii) S2, which bears the membrane-fusion elements. Similar to S_CoV-1_, the shedding of S1 renders accessible on S2 the second proteolytic cleavage site 797^R^, namely S2’ (Belouzard et al., 2009). Depending on the cell or tissue type, one or several host proteases, including furin, trypsin, cathepsins or transmembrane protease serine protease-2 or −4, can be involved in this second cleavage step (Coutard et al., 2020). The consequent “fusogenic” conformational changes of S result in the exposure of a Fusion Peptide (FP), adjacent to S2’. Insertion of FP to the host cell/vesicle membrane primes the fusion reaction, which in turn leads to the viral RNA release into the host cytosol (Lai et al., 2017). The facts that the S_CoV-2_-ACE2 interaction is the only mechanism thus far identified for the host cell infection by SARS-CoV-2, and that the RBD contains numerous conformational B-cell epitopes (Walls et al., 2020), designate this viral envelope glycoprotein as the main target for neutralizing antibodies (NAbs).

Compared to: (i) attenuated or inactivated viral vaccine candidates which require extensive safety testing, (ii) nucleic acid with moderate immunogenicity in human linked to the difficulty of their delivery to targeted immune cells (Hobernik and Bros, 2018), (iii) protein vaccines which require the use of adjuvants and boosting, viral vector vaccines such as adenoviral vectors are interesting vaccine candidates that generate strong immune responses. However, adenoviral vectors are target of pre-existing immunity in the human population, which largely reduces their immunogenicity (Rosenberg et al., 1998; Schirmbeck et al., 2008). Non-replicative lentiviral vaccinal vectors (LV) possess a large potential at eliciting strong and long-lasting adaptive immunity, including Ab responses (Di Nunzio et al., 2012; Hu et al., 2011; Ku et al., 2020; Zennou et al., 2000). These vectors induce very minor inflammation (Lopez et al., in preparation) and their safety has been demonstrated in human in a phase 1 HIV-1 vaccine trial (2011-006260-52 EN). In addition, LV are pseudo-typed with the envelope glycoprotein of Vesicular Stomatitis Virus (VSV-G), to which human population is barely exposed. This minimizes the risk of vaccine efficacy reduction linked to a pre-existing cross-reactive immunity (Hu et al., 2011).

To develop a vaccine candidate able to induce NAbs specific to S_CoV-2_, we generated LV coding for: (i) S1 alone (LV::S1), (ii) S1-S2 ectodomain, without the transmembrane and internal tail domains (LV::S1-S2), or (iii) full-length, membrane anchored form of S (LV::S_FL_). We established that LV::S_FL_ gave rise to elevated amounts of NAbs, which inhibit ACE2^+^ host-cell invasion by S_CoV-2_-pseudo-typed virions. Anti-S_CoV-2_ CD8^+^ T cell effectors were also efficiently induced in LV::S_FL_-immunized mice. Moreover, in a mouse model in which the expression of human ACE2 (hACE2) was induced in the respiratory tracts by instillation with an adenoviral vector serotype 5 (Ad5), as well as in SARS-CoV-2-susceptible golden hamsters, we demonstrated a strong prophylactic effect of LV::S_FL_ immunization against the replication of a SARS-CoV-2 clinical isolate, accompanied by the reduction of infection-related inflammation in the lungs. Importantly, boost/target immunization with LV::S_FL_ via nasal route was instrumental in the protection efficacy. Our virological, immunological and histopathological criteria in two preclinical animal models provided the proof-of-principle evidence of marked prophylactic effects of LV-based vaccine strategies against SARS-CoV-2, and requirement for mucosal immunization to reach vigorous protective lung immunity against COVID-19.

## Results

### Induction of antibody responses by LV encoding SARS-CoV-2 Spike protein variants

To develop a vaccine candidate able to induce NAbs against S_CoV-2_, we generated LV harboring, under the transcriptional control of the cytomegalovirus (CMV) immediate-early promoter, codon-optimized sequences encoding for: (i) S1 alone (LV::S1), (ii) S1-S2 ectodomain, without the transmembrane and C-terminal short internal tail (LV::S1-S2), and (iii) full-length, membrane anchored form of S (LV::S_FL_), which all harbor the RBD (Figure 1A, Figure S1), albeit with potential conformational heterogeneities (Yuan et al., 2020). To evaluate the humoral responses induced by these vectors, C57BL/6 mice (*n* = 4/group) were immunized by a single intraperitoneal (i.p.) injection of 1 × 10^7^ Transduction Units (TU) of either LV, or an LV encoding GFP as negative control. Anti-S_CoV-2_ Ab responses were investigated in the sera at weeks 1, 2, 3, 4 and 6 post immunization (Figure 1B). In LV::S_FL_ or LV::S1-S2-immunized mice, anti-S_CoV-2_ immunoglobulin G (IgG) were detectable as early as 1 week post immunization and increased progressively until week 6 post immunization, achieving mean titer (1/dilution) ± SEM of (4.5 ± 2.9) × 10^6^ or (1.5 ± 1) × 10^6^, respectively. In comparison, anti-S_CoV-2_ IgG titers were 2 orders of magnitude lower, i.e., (7.1 ± 6.1) ×10^4^, in their LV::S1-immunized counterparts.

**Figure 1.**
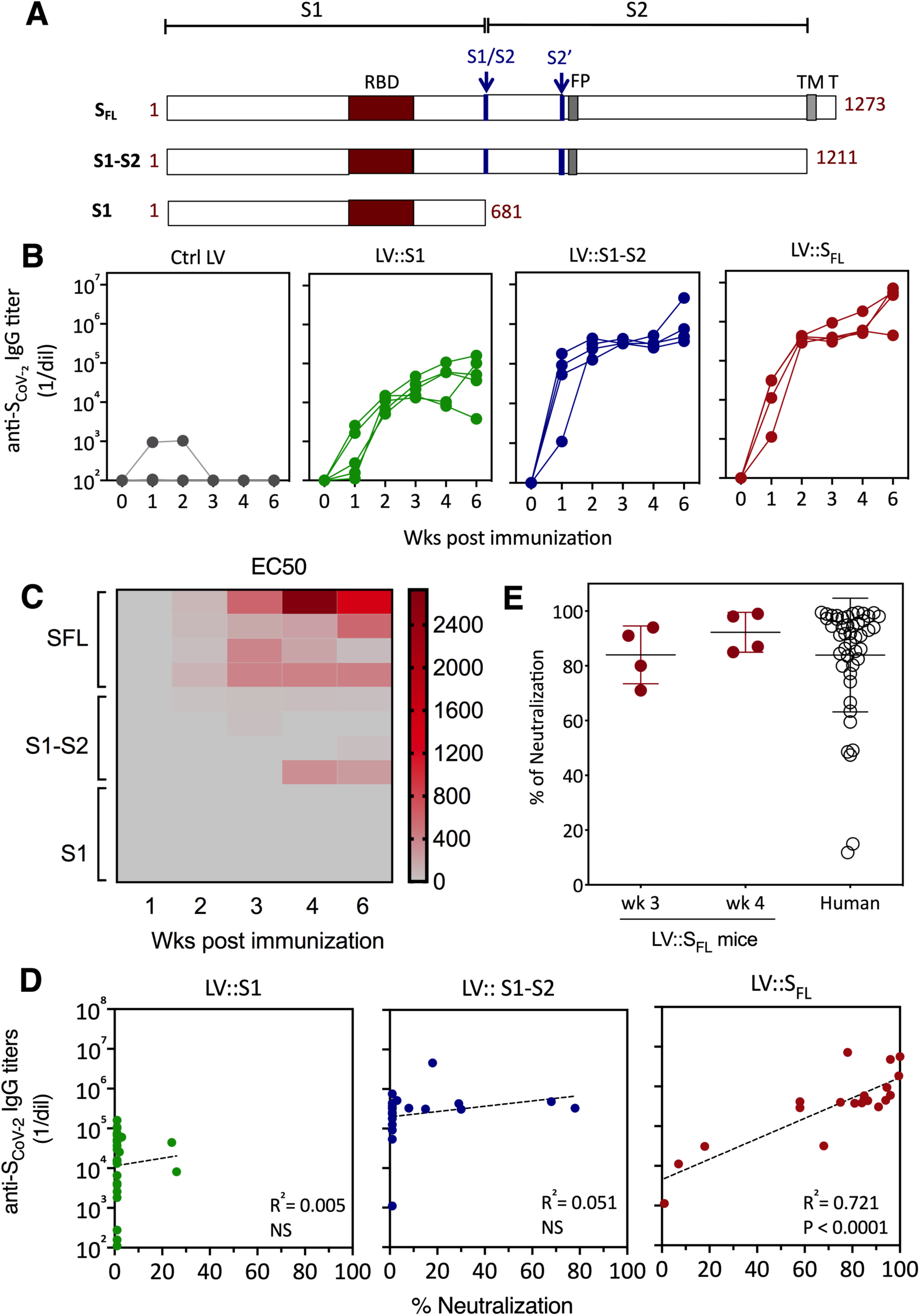
Induction of anti-S_CoV-2_ Ab responses by LV. **(A)** Schematic representation of 3 forms of S_CoV-2_ protein (S_FL_, S1-S2 and S1) encoded by LV injected to mice. RBD, S1/S2 and S2’ cleavage sites, Fusion Peptide (FP), TransMembrane (TM) and short internal tail (T) are indicated. **(B)** Dynamic of anti-S_CoV-2_ Ab response following LV immunization. C57BL/6 mice (*n* = 4/group) were injected i.p. with 1 × 10^7^ TU of LV::GFP as a negative control, LV::S1, LV::S1-S2, or LV::S_FL_. Sera were collected at 2, 3, 4 and 6 weeks post immunization. Anti-S_CoV-2_ IgG responses were evaluated by ELISA and expressed as mean endpoint dilution titers. **(C)** Neutralization capacity of anti-S_CoV-2_ Abs induced by LV::S_FL_ immunization. Mouse sera were evaluated in a sero-neutralization assay to determine 50% effective concentration (EC50) neutralizing titers. **(D)** Correlation between the Ab titers and neutralization activity in various experimental groups. Statistical significance was determined by two-sided Spearman rank-correlation test. NS: not significant. **(E)** Head-to-head comparison at a 1:40 dilution between mouse sera taken at weeks 3 or 4 after immunization and a cohort of mildly symptomatic individuals living in Crépy-en-Valois, Ile de France. These patients did not seek medical attention and recovered from COVID-19. Results are expressed as mean ± SEM percentages of inhibition of luciferase activity. See also Figure S1.

Sera were then evaluated for their capacity to neutralize SARS-CoV-2, using a reliable neutralization assay based on NAb-mediated inhibition of hACE2^+^ cell invasion by non-replicative LV particle surrogates, pseudo-typed with S_CoV-2_ (Sterlin et al., 2020). Such S_CoV-2_ pseudo-typed LV particles, harbor a luciferase reporter gene, which allows quantitation of the hACE2^+^ host cell invasion. In this assay, the luciferase activity is inversely proportional to the neutralization efficiency of NAbs present in biological fluids. Fine comparison, by 50% Effective Concentrations (EC50) assays of the sera from the LV::S1-, LV::S1-S2- or LV::S_FL_-immunized mice clearly established that LV::S_FL_ was the most potent vector at inducing anti-S_CoV-2_ NAbs (Figure 1C). Moreover, neutralization activities were correlated with anti-S_CoV-2_ IgG titers only in the sera of LV::S_FL_-immunized mice (Figure 1D). These results suggest that in the S1-S2 or S1 polypeptides, the conformations of relevant B-cell epitopes are distinct from those of the native S_FL_ and that the blocking action of NAbs is linked to their specific recognition of the native conformational epitopes. Comparison of the sera from the LV::S_FL_-immunized mice and a cohort of mildly symptomatic infected individuals living in Crépy-en-Valois, one of the first epidemic zones to appear in France, revealed similar mean neutralizing activities (Figure 1E). These data predict a potentially protective humoral response induced by LV::S_FL_.

LV::S_FL_-immunized C57BL/6 mice (*n* = 3) also displayed strong anti-S_CoV-2_ T-cell responses, as detected at week 2 post immunization by IFNγ ELISPOT-based epitope mapping, applied to splenocytes stimulated with distinct pools of 15-mer peptides spanning the full-length S_CoV-2_ (Figure 2A). Significant amounts of responding T cells were detected for 6 out of 16 peptide pools. Deconvolution of these positive pools allowed identification of S:256-275 (SGWTAGAAAYYVGYLQPRTF), S:536-550 (NKCVNFNFNGLTGTG) and S:576:590 (VRDPQTLEILDITPC) immunodominant epitopes, giving rise to > 2000 Spot Forming Unit (SFU) / spleen (Figure 2B). These epitopes elicited CD8^+^ - but not CD4^+^ - T cells, as assessed by intracellular cytokine staining (Figure 2C). The predominant CD8^+^ phenotype of these T cells is in accordance with the favored orientation of LV-encoded antigens to the MHC-I presentation pathway (Hu et al., 2011). We also identified S:441-455 (LDSKVGGNYNYLYRL), S:671-685 (CASYQTQTNSPRRAR) and S:991-1005 (VQIDRLITGRLQSLQ) subdominant epitopes, which gave rise to < 2000 SFU / spleen in ELISPOT assay (Figure 2B).

**Figure 2.**
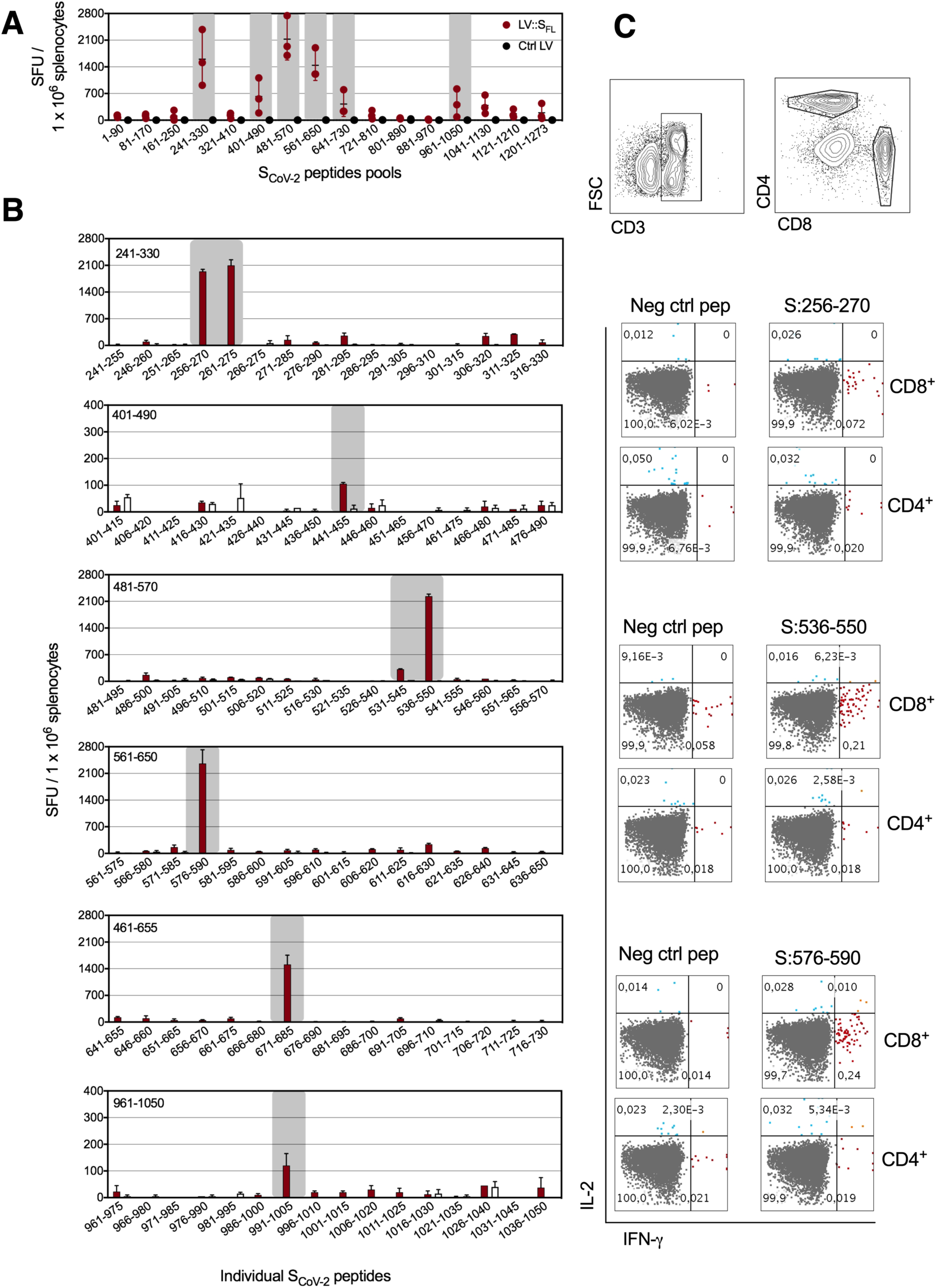
Induction of T-cell responses by LV::S_FL_. C57BL/6 mice (*n* = 3) were immunized i.p. with 1 × 10^7^ TU of LV::S_FL_ or a negative control LV. **(A)** Splenocytes collected 2 weeks after immunization were subjected to an IFN-γ ELISPOT using 16 distinct pools of 15-mer peptides spanning the entire S_CoV-2_ (1-1273 a.a.) and overlapping each other by 10 a.a. residues. SFU = Spot-Forming Cells. **(B)** Deconvolution of the 16 positive peptide pools by ELISPOT applied to splenocytes pooled from 3 LV::S_FL_-or Ctrl LV-immunized mice. **(C)** Intracellular IFN-γ *versus* IL-2 staining of CD4^+^ or CD8^+^ T splenocytes after stimulation with individual peptides encompassing the immuodominant epitopes.

### Establishment of a murine model expressing hACE2 in the respiratory tracts

As S_CoV-2_ does not interact well with murine ACE2, wild-type laboratory mice are not permissive to replication of SARS-CoV-2 clinical isolates. Therefore, we sought to develop a murine model in which human ACE2 (hACE2) expression was induced in the respiratory tracts and pulmonary mucosa to evaluate the LV::S_FL_ vaccine efficacy. This method has been successfully used to establish the expression of human DPP4 for the study of mouse infection with MERS-CoV (Zhao et al., 2014) and also for hACE2 during the preparation of this manuscript (Sun et al., 2020). We generated an Ad5 vector to deliver the gene encoding for hACE2 under the transcriptional control of the CMV promoter (Ad5::hACE2) in an episomal form. We first checked in vitro the potential of the Ad5::hACE2 vector to transduce HEK293T cells by Reverse Transcriptase (RT)-PCR (Figure 3A). To achieve in vivo transduction of respiratory tract cells, we instilled intranasally (i.n.) 2.5 × 10^9^ Infectious Genome Units (IGU) of Ad5::hACE2 into C57BL/6 mice. Four days later, the hACE2 protein expression was detectable in the lung cell homogenate by Western Blot (Figure 3B). To get more insights into the in vivo expression profile of a transgene administered under these conditions, we instilled i.n. the same dose of an Ad5::GFP reporter vector into C57BL/6 mice. As evaluated by flow cytometry, 4 days post instillation, the GFP reporter was expressed not only in the lung epithelial EpCam^+^ cells, but also in lung immune cells, as tracked by the CD45 pan-hematopoietic marker (Figure 3C), showing that through this approach the transduction was efficiently achieved in epithelial cells, although not restricted to these cells.

**Figure 3.**
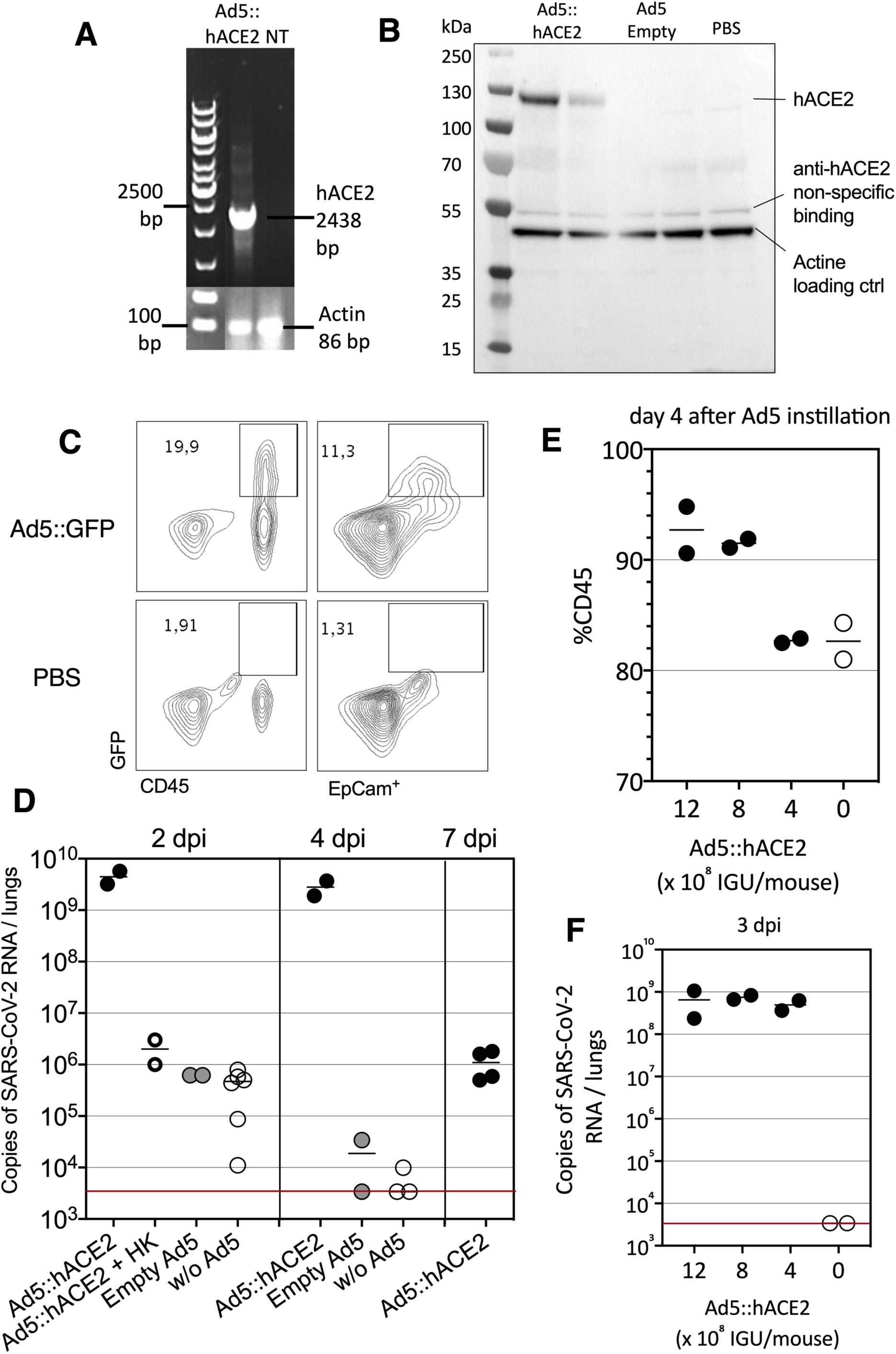
Set up of a murine model expressing hACE2 in the respiratory tracts. **(A)** Detection of hACE2 expression by RT-PCR in HEK293 T cells transduced with Ad5::hACE2, at 2 days post transduction. NT: Not transduced. **(B)** hACE2 protein detection by Western Blot in lung cell extracts recovered at day 4 after i.n. instillation of Ad5::hACE2 or empty Ad5 to C57BL/6 mice (*n* = 2/group). **(C)** GFP expression in lung cells prepared at day 4 after i.n. instillation of Ad5::GFP or PBS into C57BL/6 mice, as assessed by flow cytometry in the CD45^+^ hematopoietic or EpCam^+^ epithelial cells. **(D)** Lung viral loads in mice pretreated with 2.5 × 10^9^ IGU of Ad5::hACE2, control empty Ad5 or PBS followed by i.n. inoculation of 1 × 10^5^ TCID_50_ of SARS-CoV-2 4 days later. In one group, the Ad5::hACE2-pretreated mice were inoculated with an equivalent amounts of heat-killed (HK) virus to measure the input viral RNA in the absence of viral replication. Viral load quantitation by qRT-PCR in the lung homogenates at 2, 4 or 7 dpi. The red line indicates the detection limit. **(E)** Percentages of CD45^+^ cells in the lungs, as determined 4 days after pretreatment with various doses of Ad5::hACE2. **(F)** Lung viral loads in mice pretreated with various doses of Ad5::hACE2, followed by i.n. inoculation of 1 × 10^5^ TCID_50_ of SARS-CoV-2 4 days later. Viral load were determined at 3 dpi. See also Figure S2.

To evaluate the permissibility of such hACE2-transduced mice to SARS-CoV-2 infection, 4 days after i.n. pretreatment with either Ad5::hACE2 or an empty Ad5 control vector, C57BL/6 mice were inoculated i.n. with 1 × 10^5^ TCID_50_ (Median Tissue Culture Infectious Dose) of a SARS-CoV-2 clinical isolate (BetaCoV/France/IDF0372/2020), which was isolated in January 2020 from a COVID-19 patient by the National Reference Centre for Respiratory Viruses (Institut Pasteur, Paris, France) (Lescure et al., 2020). The lung viral loads, determined at 2 days post inoculation (dpi) by reverse transcription and quantitative real-time PCR (qRT-PCR), were as high as (4.4 ± 1.8) × 10^9^ copies of SARS-CoV-2 RNA in Ad5::hACE2-pretreated mice, compared to only (6.2 ± 0.5) × 10^5^ copies in empty Ad5-pretreated mice, or (4.0 ± 2.9) × 10^5^ copies in un-pretreated mice (Figure 3D). In the latter two control groups, these copy numbers corresponded to the input viral RNA, as determined in Ad5::hACE2-pretreated mice, inoculated with equivalent amounts of heat-killed viral particles (Figure 3D). At 4 dpi, the lung viral loads were maintained in Ad5::hACE2-pretreated mice (2.8 ± 1.3 × 10^9^ copies), whereas a drop to (1.7 ± 2.3) × 10^4^ or (3.9 ± 5.1) × 10^3^ copies was observed in empty Ad5-pretreated or non-pretreated mice, respectively. In Ad5::hACE2-pretreated mice, the viral loads were still detectable at 7 dpi ((1.33 ± 0.9) × 10^6^ copies).

Ad5::hACE-2 i.n. instillation induced CD45^+^ cell recruitment to the lungs (Figure 3E). However, no pro-inflammatory effect was seen with a lower dose of 4 × 10^8^ IGU/mouse (Figure 3E), which still conferred full permissibility to SARS-CoV-2 replication (Figure 3F), and this dose was therefore chosen for the subsequent experiments described below. In this model of permissive mice, SARS-CoV-2 infection resulted in widespread infiltration of the lung interstitium by mononuclear inflammatory cells, i.e., lymphocytes and macrophages, at 3 dpi (Figure S2).

These results show that pretreatment of mice with appropriate doses of Ad5::hACE2 can render mice permissive to SARS-CoV-2 replication without inducing Ad5-mediated inflammation, thus providing a valuable model for vaccine or drug studies.

### Intranasal boost with LV::S_FL_ protects strongly against SARS-CoV-2 in mice

To investigate the prophylactic potential of LV::S_FL_ against SARS-CoV-2, C57BL/6 mice (*n* = 4-5/group) were injected i.p. with a single dose of 1 × 10^7^ TU of LV::S_FL_ or a negative control LV (sham). At week 7 post immunization, mice were pretreated with Ad5::hACE2, and 4 days later, inoculated i.n. with 1 × 10^5^ TCID_50_ of SARS-CoV-2 (Figure S3A). At 3 dpi, the lung viral loads in LV::S_FL_-vaccinated mice were reduced by ∼ 6.5 folds, i.e., mean ± SD of (5.5 ± 3.8) × 10^8^ SARS-CoV-2 RNA copies compared to (3.1 ± 1.9) × 10^9^ or (4.3 ± 3.0) × 10^9^ copies in the un- or sham-vaccinated mice, respectively (Figure S3B). Therefore, a single i.p. LV::S_FL_ injection provided partial protection in the lung, despite intense serum NAb activity.

To further improve the prophylactic effect, we evaluated the prime-boost or prime-target approaches. C57BL/6 mice (*n* = 4-5/group) were primed i.p. with 1 × 10^7^ TU of LV::S_FL_ or a control LV at week 0, and then boosted at week 3 with: (i) 1 × 10^7^ TU of the same LV via the i.p. route (“LV::S_FL_ i.p.-i.p.”, prime-boost), or (ii) with 3 × 10^7^ TU via the i.n. route (“LV::S_FL_ i.p.-i.n.”, prime-target) to attract the mediators of systemic immunity to the lung mucosa (Figure 4A). Systemic boosting with LV::S_FL_ via i.p. resulted in a significant increase in the anti-S_CoV-2_ IgG titers, which was more obvious when the binding was evaluated against the foldon-trimerized full-length S (Figure 4B, left) than against the S1 or RBD fragments (Figure S4A). This observation may suggest that the concerned B-cell epitopes are of conformational type. In contrast, mucosal targeting with LV::S_FL_ via i.n. did not lead to a statistically significant improvement of anti-S_CoV-2_ IgG titers at the systemic level (Figure 4B left, Figure S4A). In terms of serum neutralization potential, even though a trend to increase was observed after i.p. or i.n. boost, the differences did not reach statistical significance (Figure 4B right).

**Figure 4.**
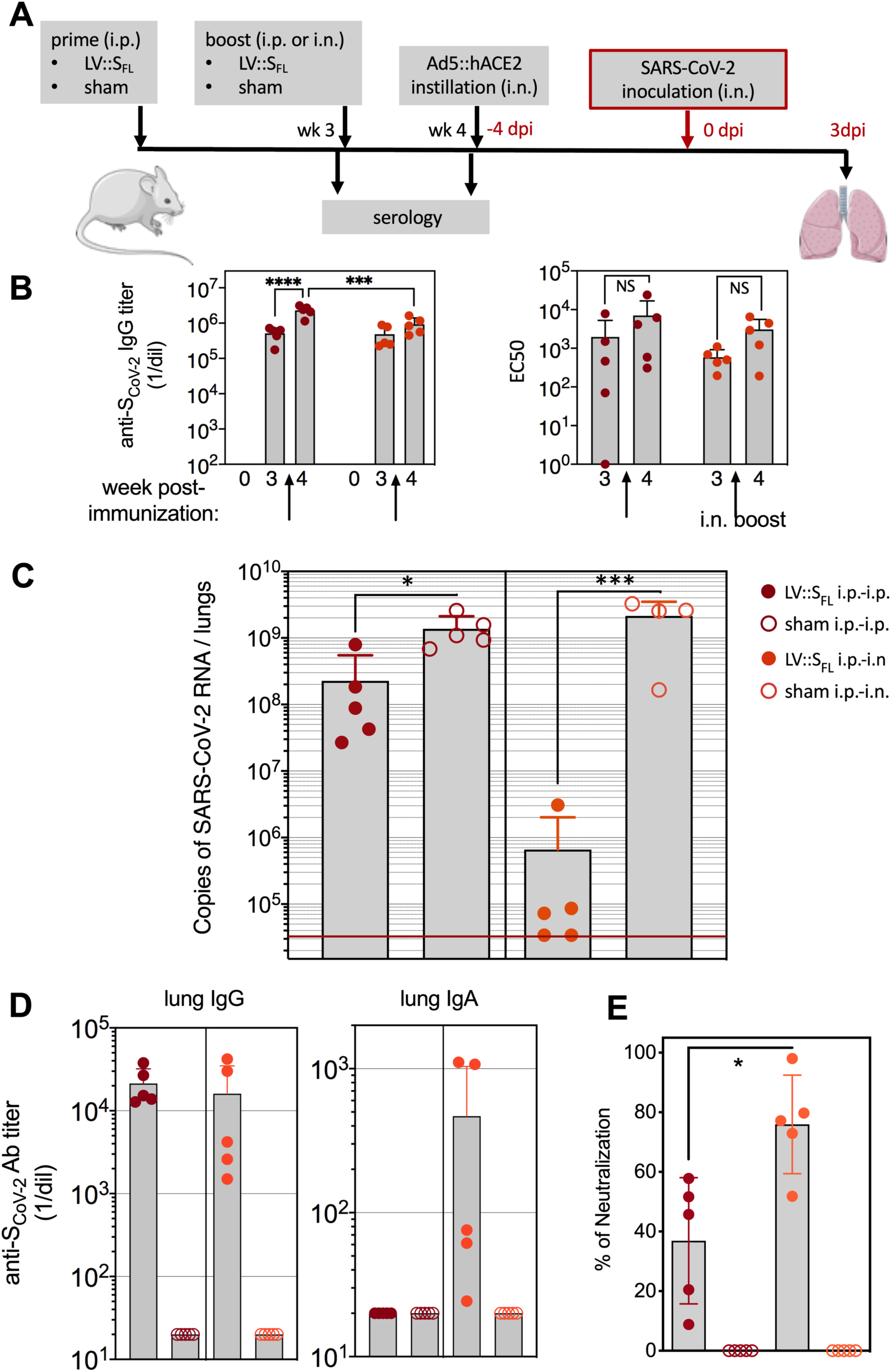
Intranasal boost with LV::S_FL_ strongly protects against SARS-CoV-2 in mice. **(A)** Timeline of the prime-boost strategy based on LV, followed by Ad5::hACE2 pretreatment and SARS-CoV-2 challenge. **(B)** Titers of anti-S_CoV-2_ IgG, as quantitated by ELISA in the sera of C57BL/6 mice primed i.p. at week 0 and boosted i.p. or i.n. at week 3 (left). Titers were determined as mean endpoint dilution before boost (week 3) and challenge (week 4). *** *p* <0.001, **** *p* <0.0001; two-way ANOVA followed by Sidak’s multiple comparison test. NS, not significant. Neutralization capacity of these sera, indicated as EC50 (right). See also Figure S4A. **(C)**. Lung viral loads at 3 dpi in mice primed (i.p.) and boosted (i.p. or i.n.) with LV::S_FL_. Sham-vaccinated received an empty LV. The red line indicates the detection limit. Statistical significance of the differences in the viral loads was evaluated by two tailed unpaired t test; * = *p* <0.0139, *** = *p* <0.0088. **(D)** Titers of anti-S_CoV-2_ IgG and IgA Abs determined in the clarified lung homogenates by ELISA, by use of a foldon-trimerized S_CoV-2_ for coating. See also Figure S4B. **(E)** Neutralizing activity of the clarified lung homogenates, determined for 1/5 dilution. Statistical significance of the difference was evaluated by Mann-Whitney U test (*= *p* <0.0159).

All mice were then pretreated with Ad5::hACE2 and challenged i.n. with 0.3 × 10^5^ TCID_50_ of SARS-CoV-2 at week 4 post prime. At 3 dpi, the lung viral loads were significantly lower in LV::S_FL_ i.p.-i.p. immunized mice, i.e., mean ± SD (2.3 ± 3.2) × 10^8^, than in sham-vaccinated mice (13.7 ± 7.5) × 10^8^ copies of SARS-CoV-2 RNA, (Figure 4C). This viral load reduction was similar to that obtained with a single LV::S_FL_ administration (Figure S3B). Most importantly, after i.n. LV::S_FL_ target immunization, > 3 log10 decrease in viral loads was observed and 2 out of 5 mice harbored undetectable lung viral loads as determined by qRT-PCR assay. Anti-S_CoV-2_ IgG were detected in the clarified lung homogenates of the partially (LV::S_FL_ i.p.-i.p.) or the fully (LV::S_FL_ i.p.-i.n.) protected mice. In contrast anti-S_CoV-2_ IgA were only detectable in the fully protected LV::S_FL_ i.p.-i.n. mice (Figure 4D). Higher neutralizing activity was detected in the clarified lung homogenates of LV::S_FL_ i.p.-i.n. mice than of their LV::S_FL_ i.p.-i.p. counterparts (Figure 4E). Therefore, increasing the titers of NAb of IgG isotype at the systemic levels did not improve the protection against SARS-CoV-2. However, a mucosal i.n. target immunization, with the potential to attract immune effectors to the infection site and able to induce local IgA Abs, correlated with the inhibition of SARS-CoV-2 replication.

Because of the importance of innate immune hyperactivity in the pathophysiology of COVID-19 (Vabret et al., 2020), we investigated the lung innate immune cell subsets in the non-infected controls, sham-vaccinated or LV::S_FL_-vaccinated mice inoculated with SARS-CoV-2 (Figure 5A). At 3 dpi, we detected no difference in the proportions of basophils or NK cells versus total lung CD45^+^ cells among the experimental groups (Figure 5B). In sharp contrast, we detected higher proportions of alveolar macrophages, dendritic cells, mast cells, Ly6C^+^ or Ly6C^-^ monocytes/macrophages or neutrophils versus total lung CD45^+^ cells, in sham-vaccinated mice. These observations demonstrate that in this mouse model, lung SARS-CoV-2 loads correlate with the expansion of several inflammation-related innate immune cell subsets, while vaccine-mediated protection dampens or prevents the inflammatory reaction.

**Figure 5.**
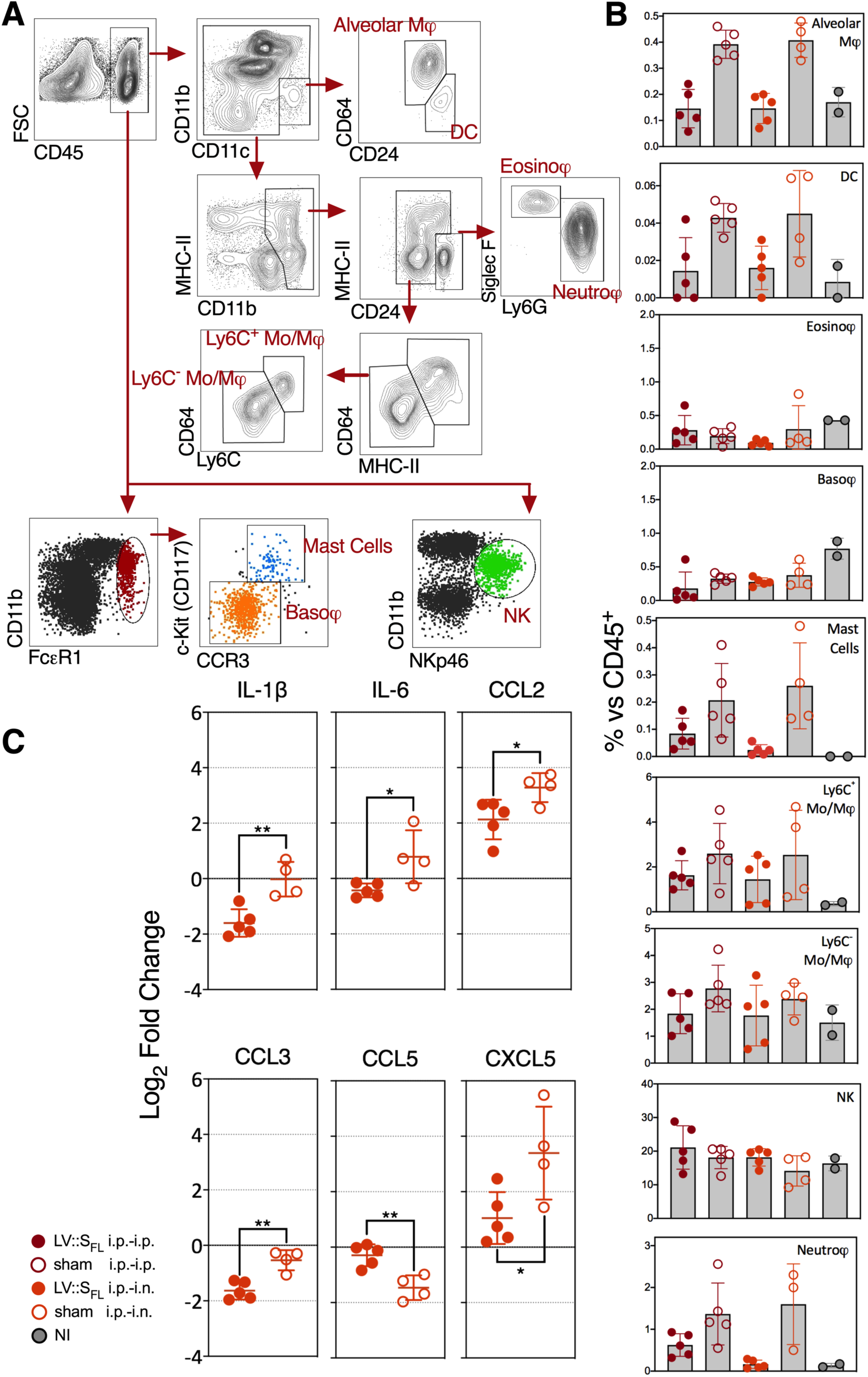
LV::S_FL_ vaccination reduces SARS-Co-2-mediated lung inflammation in mice. **(A)** Gating strategy applied to total lung cells to quantitate innate immune subsets. **(B)** Percentages of each innate immune subset versus total lung CD45^+^ cells at 3 dpi in mice sham-vaccinated or vaccinated with LV::S_FL_, following various prime-boost regimen compared to non-infected (NI) controls which only received PBS. All mice were pretreated with Ad5::hACE-2, 4 days prior to SARS-CoV-2 inoculation. **(C)** Relative log_2_ fold change in cytokines and chemokines mRNA expression in mice sham-vaccinated or vaccinated with LV::S_FL_, following various prime-boost regimen at 3 dpi. Data were normalized versus PBS-treated, unchallenged controls. Statistical significance of the differences in cytokines and chemokines level was evaluated by one-way ANOVA; * = *p*<0.05, ** = *p* <0.01. See also Figure S5A-C.

This was corroborated by the reduced inflammatory cytokine and chemokine contents in the RNA extracted from total lung homogenates of mice immunized with “LV::S_FL_ i.p.-i.n.” (Figure 5C, Figure S5A) or “LV::S_FL_ i.p.-i.p.” (Figure S5B) and protected against SARS-CoV-2. Significant reductions were notably detected in the IL-1β, IL-6, CCL2, CCL3 and CXCL5 contents in the lungs of “LV::S_FL_ i.p.-i.n.”- versus sham i.p.-i.n.-vaccinated and challenged mice (Figure 5C, Figure S5A). Significant reductions of TNF-α, TGF-β, IL-1β, IL-12p40, IL-17A, IL-33, CCL2, CCL3, CCL5, CXCL9 and CXCL10 contents were also observed in the lungs of “LV::S_FL_ i.p.-i.p.”- versus sham i.p.-i.p.- vaccinated and challenged mice (Figure S5B). The expression of the other mediators tested was not significantly changed (Figure S5A, C). Therefore, the conferred protection also avoided pulmonary inflammation mediated by SARS-CoV-2 infection, as demonstrated by cytometric or qRT-PCR approaches.

### Intranasal vaccination with LV::S_FL_ strongly protects against SARS-CoV-2 in golden hamsters

Outbred *Mesocricetus auratus*, so-called golden hamsters, provide a suitable pre-clinical model to study the COVID-19 pathology, since the ACE2 ortholog of this species interacts productively with S_CoV-2_, supporting host cell invasion and viral replication (Sia et al., 2020). We thus investigated in this model the protective effect against SARS-CoV-2 infection of vaccination by LV::S_FL_ and by an integrase deficient, non-integrative version of this vector (NILV), with the prospect of application in future clinical trials.

To assess the prophylactic effect of vaccination following prime-boost/target regimen, *M. auratus* hamsters (*n* = 6/group) were: (i) primed i.p. with 1 × 10^6^ TU of LV::S_FL_ and boosted i.n. at week 5 with 4 × 10^7^ TU of LV::S_FL_, (“int LV::S_FL_ i.p.-i.n. Low”), (ii) primed i.p. with 1 × 10^7^ TU of LV::S_FL_ and boosted i.n. at week 5 with 4 × 10^7^ TU of LV::S_FL_ (“int LV::S_FL_ i.p.-i.n. High”), or (iii) primed intramuscularly (i.m.) with 1 × 10^8^ TU of NILV::S_FL_ and boosted i.n. at week 5 with 1 × 10^8^ TU of NILV::S_FL_ (“NILV::S_FL_ i.m.-i.n.”) (Figure 6A). Sham-vaccinated controls received the same amounts of an empty LV via i.p. and then i.n. routes. Strong and comparable anti-S_CoV-2_ IgG Abs were detected by ELISA in the sera of hamsters from the three vaccinated groups, before and after the i.n. boost (Figure 6B). Post boost/target serology detected neutralization activity in all groups, with the highest EC50 average observed in “int LV::S_FL_ i.p.-i.n. High” individuals. Such levels were comparable to those detected in COVID-19 cases or healthy contacts in humans (Figure 6C). All hamsters were challenged i.n. with 0.3 × 10^5^ TCID_50_ of SARS-CoV-2 at week 5. Up to 16% weight loss was progressively reached at 4 dpi in sham-vaccinated individuals, compared to a non-significant loss in the LV::S_FL_-vaccinated groups (Figure 6D). As assessed by qRT-PCR, at 2 dpi, decreases of ∼ 1.2-to-2 log10 were observed in the lung viral loads of “int LV::S_FL_ i.p.-i.n. Low”, “int LV::S_FL_ i.p.-i.n. High” and “NILV::S_FL_ i.m.-i.n.” groups, compared to sham-vaccinated hamsters (Figure 6E). At 4 dpi, the magnitude of viral load reductions in the vaccinated groups were much more important and reached >3 log10, compared to the sham-vaccinated individuals. Using the Plaque Forming Unit (PFU) assay, at 2 and 4 dpi, we detected (6.56 ± 1.88) and (2.94 ± 3.16) × 10^6^ PFU/lung of sham-vaccinated controls. At 2 dpi, the decrease in the viral loads were largely statistically significant in all of the LV::S_FL_- vaccinated hamsters and at 4 dpi, with the exception of one “NILV::S_FL_ i.m.-i.n.” hamster, no vaccinated individuals possessed detectable infectious viral particles (Figure 6F).

**Figure 6.**
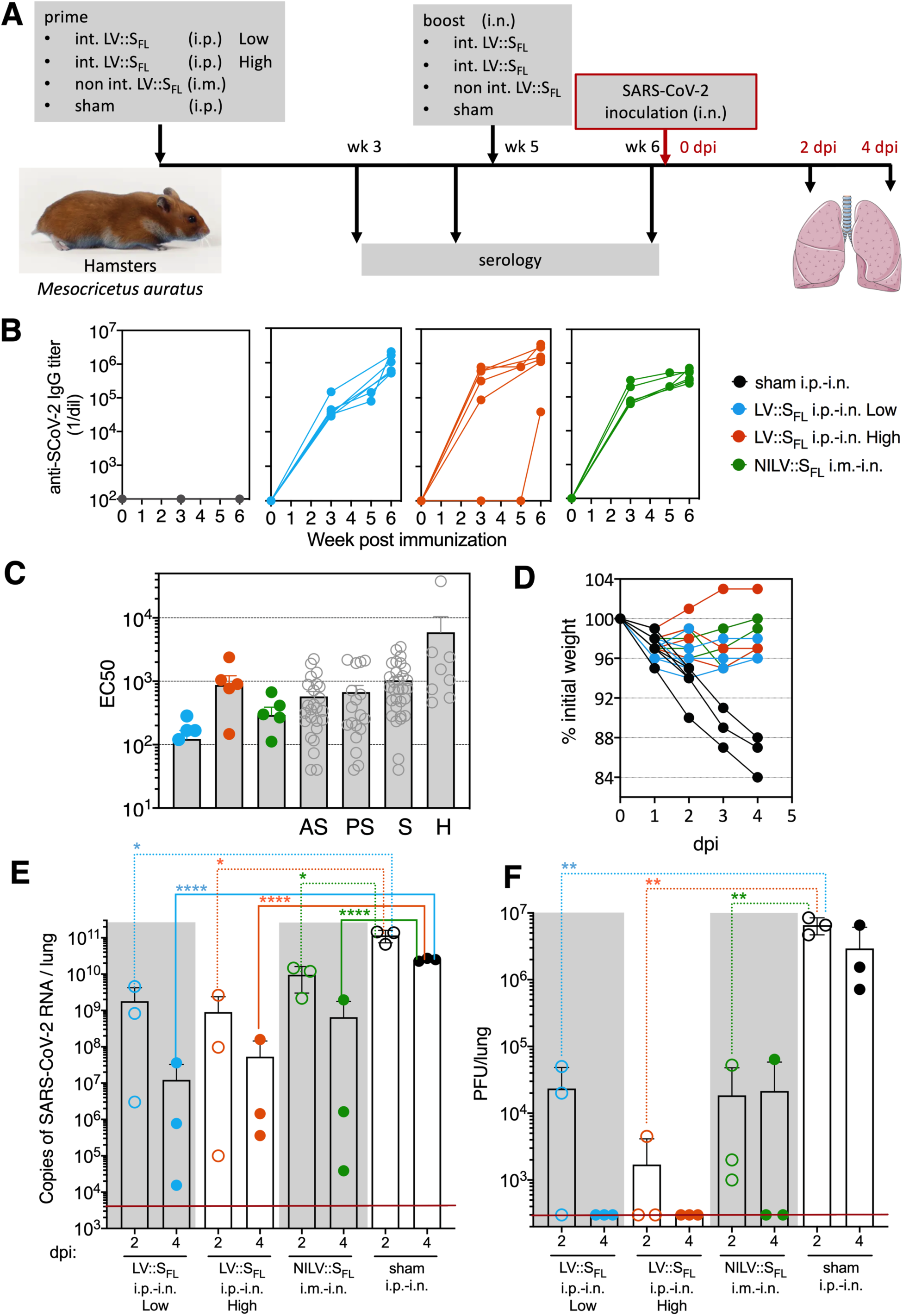
Intranasal vaccination with LV::S_FL_ strongly protects against SARS-CoV-2 in golden hamsters. **(A)** Timeline of the LV::S_FL_ prime-boost/target immunization regimen and SARS-CoV-2 challenge in hamsters. Sham-vaccinated received an empty LV. **(B)** Dynamic of anti-S_CoV-2_ Ab response following LV immunization. Sera were collected from sham- or LV-vaccinated hamsters at 3, 5 (pre-boost), and 6 (post-boost) weeks after the prime injection. Anti-S_CoV-2_ IgG responses were evaluated by ELISA and expressed as mean endpoint dilution titers. **(C)** Post boost/target EC50 neutralizing titers, determined in the hamsters’ sera after boost, and as compared to the sera from a cohort of asymptomatic (AS), pauci-symptomatic (PS), symptomatic COVID-19 cases (S) or healthy contacts (H) in humans. **(D)** Weight follow-up in hamsters, either sham- or LV::S_FL_-vaccinated with diverse regimens. For further clarity, only the individuals reaching 4 dpi are shown. Those sacrificed at 2 dpi had the same mean weight as their counterparts of the same groups between 0 and 2 dpi. **(E)** Lung viral loads at 2 or 4 dpi with SARS-CoV-2 in LV::S_FL_-vaccinated hamsters. Statistical significance of the differences in the viral loads was evaluated by two tailed unpaired t test; * = *p*<0.0402, **** = *p* <0.0001. **(F)** Lung viral infectious particles, quantified as PFU at 2 or 4 dpi. The red lines indicate the detection limit of the qRT-PCR or PFU assay.

As evaluated by qRT-PCR in the total lung homogenates of the protected “int LV::S_FL_ i.p.-i.n. Low”, “int LV::S_FL_ i.p.-i.n. High” and “NILV::S_FL_ i.m.-i.n.” groups, substantial decreases were observed at 4 dpi in the expression of inflammatory IFN-γ and IL-6 cytokines, anti-inflammatory IL-10 cytokine, and CCL2, CCL3 and CXCL10 chemokines, compared to their unprotected sham-vaccinated counterparts (Figure 7A). The other inflammatory mediators tested were not significantly modified (Figure S6).

**Figure 7.**
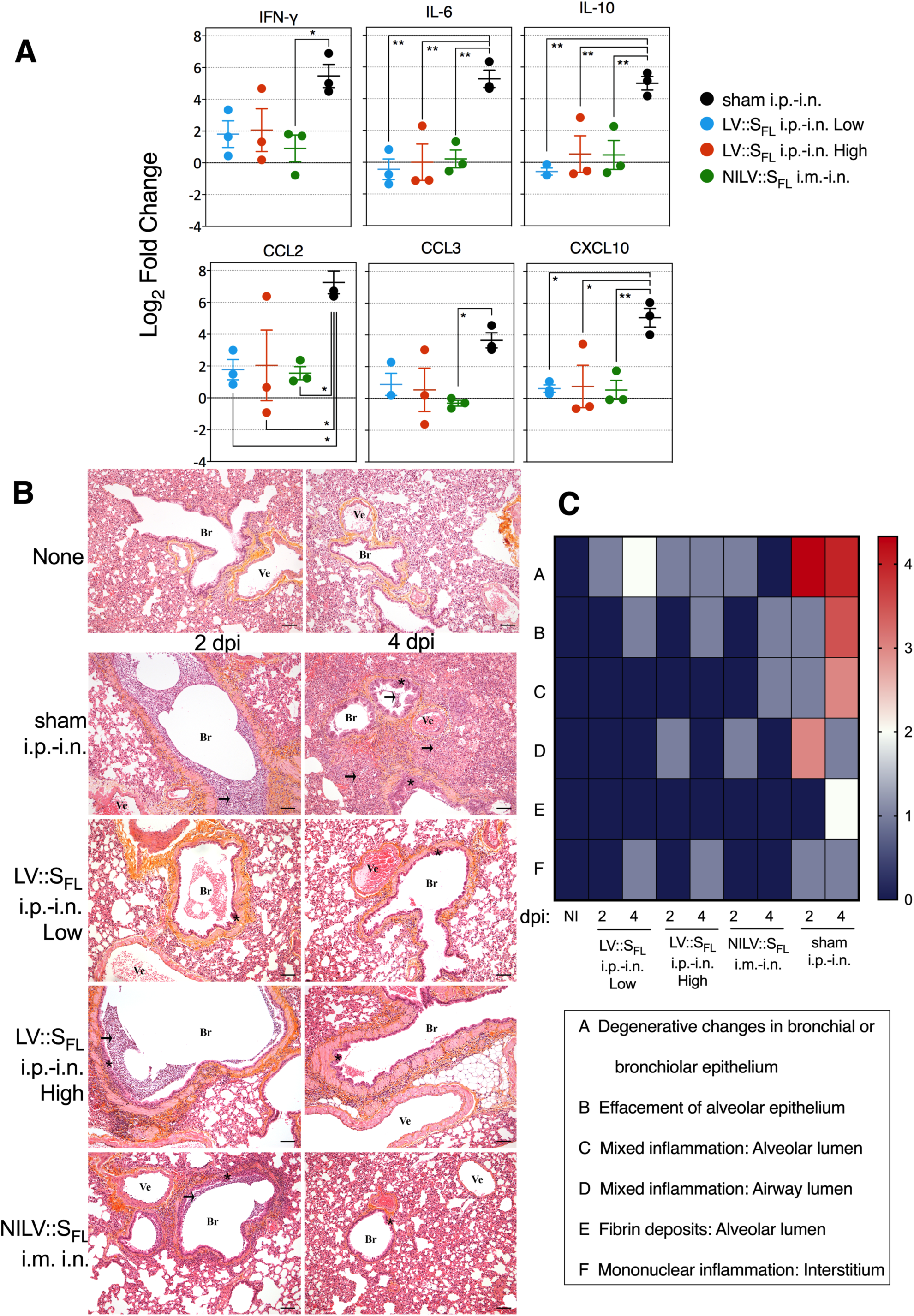
LV::S_FL_ vaccination reduces SARS-Co-2-mediated lung inflammation and histopathology in golden hamsters. Animals are those detailed in the Figure 6. **(A)** Relative log_2_ fold changes in cytokines and chemokines expression in LV::S_FL_-vaccinated and protected hamsters versus unprotected sham-vaccinated individuals, as determined at 4 dpi by qRT-PCR in the total lung homogenates and normalized versus untreated controls. Statistical significance of the differences in cytokines and chemokines level was evaluated by one-way ANOVA; * = *p*<0.05, ** = *p* <0.01. See also Figure S6. **(B)** Histological analysis HE&S lung shown for 2 and 4 dpi. Original magnification: x10, scale bar: 100 µm. Br: Bronchi or bronchiole. Bv: Blood vessel. Arrow: Mononuclear inflammatory cell infiltration. Star: Degenerative changes in the respiratory epithelium. **(C)** Heatmap recapitulating the average of histological scores, for each defined parameter and determined for individuals of the same groups at 2 or 4 dpi.

In sham-vaccinated and challenged hamsters, overall marked multifocal degenerative changes of the bronchial/bronchiolar epithelium, moderate effacement of the epithelium, associated with mild to moderate mixed inflammation in the airway lumen, and minimal multifocal interstitial mononuclear cell inflammation at both time-points (Figure 7B). At 4 dpi, effacement of the respiratory epithelium and mixed inflammation in the alveoli were more severe; moderate fibrin deposits were also noted. In vaccinated hamsters, regardless of the regardless of the vaccine dose, lung lesions were clearly of lower incidence and severity (Figure 7C). In these animals, only minimal to mild degenerative lesions and minimal effacement of the bronchial/bronchiolar epithelium and occasionally mixed inflammation in the airway lumen of alveoli and mononuclear infiltrates in the interstitium were observed. These microscopic findings were similar after vaccination with the int LV or NILV.

Altogether, based on a complete set of virological, immunological and histopathological data, the LV::S_FL_ vector elicited anti-S_CoV-2_ NAbs and T-cell responses and provided robust protection against SARS-CoV-2 infection in two pertinent animal models, and particularly when mucosal i.n. administration was included in the vaccinal scheme.

## Discussion

Prophylactic strategies are necessary to control SARS-CoV-2 infection which, 6 months into the pandemic, still continues to spread exponentially without signs of slowing down. It is now demonstrated that primary infection with SARS-CoV-2 in rhesus macaques leads to protective immunity against re-exposure (Chandrashekar et al., 2020). Numerous vaccine candidates, based on naked DNA (Yu et al., 2020) or mRNA, recombinant protein, replicating or non-replicating viral vectors, including adenoviral Ad5 vector (Zhu et al., 2020), or alum-adjuvanted inactivated virus (Gao et al., 2020) are under active development for COVID-19 prevention. Our immunological rationale for selecting LV to deliver gene encoding S_CoV-2_ antigen was based on its potential to induce in situ expression of heterologous genes, and on the ability of this immunization platform to elicit sustained humoral and cell-mediated responses. Unique to LV is the ability to transduce proliferating as well as non-dividing cells such as dendritic cells (Esslinger et al., 2002; Firat et al., 2002; He et al., 2005), thereby serving as a powerful vaccination strategy (Beignon et al., 2009; Buffa et al., 2006; Coutant et al., 2012; Gallinaro et al., 2018; Iglesias et al., 2006) to provoke strong and long-lasting adaptive responses (Cousin et al., 2019; Ku et al., in revision.). Notably, in net contrast to many other viral vectors, LV do not suffer from pre-existing immunity in populations, due to their pseudo-typing with the glycoprotein envelope from Vesicular Stomatitis Virus, to which humans are barely exposed. We recently demonstrated that a single injection of an LV expressing Zika envelope provides a rapid and durable sterilizing protection against Zika infection (Ku et al., 2020). Our recent comprehensive comparison of LV to the gold standard Ad5 immunization vector also documented the superiority of LV in inducing multifunctional and central memory T-cell responses in the mouse model, and its strong immunogenicity in outbred rats (Ku et al., in revision.), underlining the suitability of LV for vaccinal applications.

Because laboratory mice are not naturally susceptible permissive to SARS-CoV-2 infection, we set up an *in vivo* transduction murine model in which the hACE2 expression is induced in the respiratory tracts by an i.n. Ad5::hACE2 pretreatment prior to SARS-CoV-2 inoculation. Our system, very close to a recently published protocol (Sun et al., 2020) confirmed that mice become largely permissive to SARS-CoV-2 replication in the lungs and thus represent a model for assessment of vaccine or drug efficacy against this virus. Even though the Ad5::hACE2 model may not fully mimic the physiological ACE2 expression profile and thus may not reflect all the aspects of the pathophysiology of SARS-CoV-2 infection, it provides a pertinent model to evaluate *in vivo* the effects of anti-viral drugs, vaccine candidates, various mutations or genetic backgrounds on the SARS-CoV-2 replication. By using a low dose of Ad5::hACE2, no CD45^+^ cell recruitments were detectable at day 4 post instillation, indicative of an absence of Ad5-related inflammation before the inoculation of SARS-CoV-2.

We first evaluated the efficacy of several LV each encoding for one of the variants of S, i.e., the S1 domain alone (LV::S1), the S1-S2 ecto-domain, devoid of the transmembrane and C-terminal short internal tail (LV::S1-S2), or full-length, membrane anchored protein (LV::S_FL_). Even though a single administration of each of these LV was able to induce high anti-S_CoV-2_ Ab titers, only LV::S_FL_ induced highly functional Nabs with neutralizing activities similar to those found in a cohort of symptomatic SARS-CoV-2 patients. This finding predicted a protective potential of the humoral responses induced by the LV::S_FL_ vector. In addition, strong anti-S_CoV-2_ CD8^+^ T-cell responses were also observed in the spleen of mice as early as 2 weeks after a single LV::S_FL_ injection, as detected against numerous MHC-I-restricted immunogenic regions that we identified in C57BL/6 (H-2^b^) mice.

In the transduced mouse model which allows high rates of SARS-CoV-2 replication, vaccination by a single i.p. administration of 1 × 10^7^ TU of LV::S_FL_, 6 weeks before virus inoculation, was sufficient to inhibit the viral replication by ∼ 6 folds. Further boosting via the systemic route did not afford improved protection rate when compared to a single administration. However, systemic priming followed by a mucosal boost inhibited efficiently viral replication and prevented lung inflammation. This protection was correlated with high titers of anti-S_CoV-2_ IgG and IgA and a strong neutralization activity in sera. Additional experiments in appropriate transgenic mice or adoptive immune cell transfer approaches will be necessary to identify the immunological pathways that contribute to disease severity or protection against SARS-CoV-2. NAbs and cell-mediated immunity, very efficaciously induced with the LV-based vaccine candidate, may synergize to block infection and viral replication.

*M. auratus* golden hamsters are naturally permissive to SARS-CoV-2 replication and recapitulate the human COVID-19 physiopathology (Sia et al., 2020). In hamsters immunized with the prime-target regimen, either with integrative LV or NILV::S_FL_, we observed substantial degrees of protection against SARS-CoV-2, as judged by: (i) the drastic reduction of the lung viral loads (determined by qRT-PCR) or infectious particle clearance (determined by PFU assay), (ii) the significant reduction in the expression of inflammatory mediators in the lung, and (iii) the efficient prevention of lung tissue damage. Confirmation, in this highly sensitive species, of the protection results further supports the LV::S_FL_ vaccine, especially in its non-integrative form, as a candidate for future introduction into clinical trials.

Ab-Dependent Enhancement (ADE) of coronavirus entry to the host cells has been evoked as a potential obstacle in vaccination against coronaviruses. With DNA (Yu et al., 2020) or inactivated SARS-CoV-2 virus (Gao et al., 2020) vaccination in macaques, no immunopathological exacerbation has been observed. Long term observation even after decrease in Ab titer could be necessary to exclude such hypothesis. In the case of MERS-CoV, it has been reported that one particular RBD-specific neutralizing monoclonal Ab (Mersmab1) could mediate in vitro ADE of MERS-CoV into the host cells by mimicking the viral receptor human DPP4 and inducing conformational rearrangements of S_MERS_ (Wan et al., 2020). We believe that it is difficult to compare the polyclonal Ab response and its paratope repertoire complexity with the singular properties of a monoclonal Ab which cannot be representative of the polyclonal response induced by a vaccine. In addition, very contradictorily, results from the same team reported that a single-dose treatment with a humanized version of Mersmab1 afforded complete protection of a human transgenic mouse model from lethal MERS challenge (Qiu et al., 2016). Therefore, even with an Ab which might facilitate the cell host invasion in vitro in some conditions, not only there is no exacerbation of the infection in vivo, but also there is a notable protection.

It is noteworthy that all vaccine candidates currently under clinical development are planned to be administered via the i.m. route and will thus induce systemic - but not mucosal - immunity. Our results evidenced a strong correlation between the presence of mucosal anti-S_CoV-2_ IgA, local NAb activity in the respiratory tracts and very robust pulmonary protection. Therefore, vaccine administration via the i.n. route, which is the main entry door of SARS-CoV-2, has to be taken into the account when establishing the COVID-19 vaccination protocols, especially since this route is non-invasive and particularly suitable for mass vaccination of children and elderly people.

The fact that the current vaccine candidates do not take advantage of i.n. immunization is certainly related to the pro-inflammatory properties of most of the vaccination vectors or the requirement of encapsulation/adjuvantation of nucleic acids or inactivated vaccines, which rise safety concerns. Unlike most of the other vaccination viral vectors, LV are non-replicative, non-inflammatory, do not suffer from pre-existing immunity in human populations, and are effective under their non-integrative variant. Therefore, LV are particularly safe and favorable for use in mucosal immunization through the i.n. route, which is essential in immune protection against SARS-CoV-2. Prophylactic vaccination is the most cost-effective and efficient strategy against infectious diseases in general and against emerging coronaviruses in particular. Our results firmly established that the LV encoding for S_CoV-2_, used in a prime-target protocol is a prominent vaccine strategy against COVID-19.

## Acknowledgments

The authors are grateful to Pr Sylvie van der Werf (National Reference Centre for Respiratory Viruses hosted by Institut Pasteur, Paris, France) for providing the BetaCoV/France/IDF0372/2020 SARS-CoV-2 clinical isolate. The strain BetaCoV/France/IDF0372/2020 was supplied through the European Virus Archive goes Global (Evag) platform, a project that has received funding from the European Union’s Horizon 2020 research and innovation program under grant agreement No 653316. The authors thank Dr Cyril Planchais for the preparation of recombinant homotrimeric S, S1 and RBD proteins and Damien Batalie for excellent technical assistance in determination of viral loads. The preparation of histological sections was performed gratuitously by HISTALIM (Part of Cerba Research -Biomarkers Division, Montpellier, France).

This work is also supported by grants from Institut Pasteur, TheraVectys and Agence Nationale de la Recherche (ANR) HuMoCID. Min Wen Ku is part of the Pasteur -Paris University (PPU) International PhD Program and received funding from the Institut Carnot Pasteur Microbes & Santé, and the European Union’s Horizon 2020 research and innovation program under the Marie Sklodowska-Curie grant agreement No 665807.

## Author Contribution

Study concept and design: MWK, MB, FA, AM, NE, LM, PC, acquisition of data: MWK, MB, PA, JL, KN, BV, FN, PS, HT, ES, FA, AM, LM, construction and production of LV and technical support: PA, FM, AN, FN, CB, PS, analysis and interpretation of data: MWK, MB, PA, JL, KN, FA, AM, NE, LM, PC, histology: MLD, FG, LF, recombinant S proteins: HM, drafting of the manuscript: MWK, MB, FG, LM, PC.

## Methods

### Construction of transfer pFLAP plasmids coding for S_FL_, S1-S2, or S1 proteins

A codon-optimized full-length S (1-1273) sequence was amplified from pMK-RQ_S-2019-nCoV and inserted between BamHI and XhoI sites of pFlap-ieCMV-WPREm. Sequences encoding for S1-S2 (1-1211) or S1 (1-681) were amplified by PCR from the pFlap-ieCMV-S_FL_-WPREm plasmid and sub-cloned into pFlap-ieCMV-WPREm between the BamHI and XhoI restriction sites (Figure S1). Each of the PCR products were inserted between the native human ieCMV promoter and a mutated Woodchuck Posttranscriptional Regulatory Element (mWPRE) sequence, in which the *atg* starting codon was mutated to avoid transcription of the downstream truncated “X” protein of Woodchuck Hepatitis Virus, in order to improve the vector safety. Plasmids were amplified in *Escherichia coli* DH5a in Lysogeny Broth supplemented with 50 µg/ml of kanamycin and purified using the NucleoBond Xtra Maxi EF Kit (Macherey Nagel) and resuspended in Tris-EDTA Endotoxin-Free buffer overnight. Plasmid were quantified with a NanoDrop 2000c spectrophotometer (Thermo Ficher, Illkirch, France), aliquoted and stored at −20°C. Plasmid DNA were verified by enzymatic digestion and by sequencing the region proximal to the transgene insertion sites.

### Production and titration of LV

Non-replicative LV were produced in Human Embryonic Kidney (HEK)-293T cells, as previously detailed (Zennou et al., 2000). Briefly, lentiviral particles were produced by transient calcium phosphate co-transfection of HEK293T cells with the vector plasmid pTRIP/sE, a VSV-G Indiana envelope plasmid and an encapsidation plasmid (p8.74 or pD64V for the production of integration-proficient or integration-deficient vectors respectively). Supernatants were harvested at 48h post transfection, clarified by 6-minute centrifugation at 2500 rpm at 4°C. LV were aliquoted and stored at −80°C. Vector titers were determined by transducing 293T cells treated with aphidicolin. The titer, proportional to the efficacy of nuclear gene transfer, is determined as Transduction Unit (TU)/ml by qPCR on total lysates at day 3 post transduction, by use of forward 5’-TGG AGG AGG AGA TAT GAG GG-3’ and reverse 5’-CTG CTG CAC TAT ACC AGA CA-3’ primers, specific to pFLAP plasmid and forward 5’-TCT CCT CTG ACT TCA ACA GC-3’ and reverse 5’-CCC TGC ACT TTT TAA GAG CC-3’ primers specific to the host housekeeping gene *gadph* as previously described (Iglesias et al., 2006).

### Mouse and hamster studies

Female C57BL/6JRj mice (Janvier, Le Genest Saint Isle, France) were used between the age of 6 and 10 weeks. Male *Mesocricetus auratus* golden hamsters (Janvier, Le Genest Saint Isle, France) were purchased mature, i.e. 80-90 gr weight. At the beginning of the immunization regimen they weighed 100 to 120 gr. Experimentation on animals was performed in accordance with the European and French guidelines (Directive 86/609/CEE and Decree 87-848 of 19 October 1987) subsequent to approval by the Institut Pasteur Safety, Animal Care and Use Committee, protocol agreement delivered by local ethical committee (CETEA #DAP20007) and Ministry of High Education and Research APAFIS#24627-2020031117362508 v1. Animals were vaccinated with the indicated TU of LV and sera were collected at various time points post immunization to monitor binding and neutralization activities. Previous to i.m. or i.n. instillations, animals were anesthetized by i.p. injection of a mixture of Ketamine (Imalgene, 50 mg/kg) and Xylazine (Rompun, 50 mg/kg).

### SARS-CoV-2 inoculation

Hamsters or Ad5::hACE2-pretreated mice were anesthetized by i.p. injection of mixture Ketamine and Xylazine, transferred into a biosafety cabinet 3 where they were inoculated i.n. with 0.3 or 1 × 10^5^ TCID_50_ of the BetaCoV/France/IDF0372/2020 SARS-CoV-2 clinical isolate (Lescure et al., 2020), amplified in VeroE6 cells. The strain BetaCoV/France/IDF0372/2020 was supplied by the National Reference Centre for Respiratory Viruses hosted by Institut Pasteur (Paris, France) and headed by Pr. Sylvie van der Werf. The human sample from which strain BetaCoV/France/IDF0372/2020 was isolated has been provided by Dr. X. Lescure and Pr. Y. Yazdanpanah from the Bichat Hospital, Paris, France. The viral inoculum was contained in 20 µl for mice and in 50 µl for hamsters. Animals were then housed in an isolator in BioSafety Level 3 animal facilities of Institut Pasteur. The organs and fluids recovered from the animals infected with live SARS-CoV-2 were manipulated following the approved standard operating procedures of these facilities.

### Recombinant S_CoV-2_ proteins

Codon-optimized nucleotide fragments encoding a stabilized foldon-trimerized version of the SARS-CoV-2 S ectodomain (a.a. 1 to 1208), the S1 monomer (a.a. 16 to 681) and the RBD subdomain (amino acid 331 to 519) both preceded by a murine IgK leader peptide and followed by an 8xHis Tag were synthetized and cloned into pcDNA™3.1/Zeo^(+)^ expression vector (Thermo Fisher). Proteins were produced by transient co-transfection of exponentially growing Freestyle™ 293-F suspension cells (Thermo Fisher) using polyethylenimine (PEI)-precipitation method as previously described (Lorin and Mouquet, 2015). Recombinant S_**CoV-2**_ proteins were purified by affinity chromatography using the Ni Sepharose® Excel Resin according to manufacturer’s instructions (Thermo Fisher). Protein purity was evaluated by in-gel protein silver-staining using Pierce Silver Stain kit (Thermo Fisher) following SDS-PAGE in reducing and non-reducing conditions using NuPAGE™ 3-8% Tris-Acetate gels (Life Technologies). Purified proteins were dialyzed overnight against PBS using Slide-A-Lyzer® dialysis cassettes (10 kDa MW cut-off, Thermo Fisher). Protein concentration was determined using the NanoDrop™ One instrument (Thermo Fisher).

### ELISA

Ninety-six-well Nunc Polysorp plates (Nunc, Thermo Ficher) were coated overnight at 4 °C with 100 ng/well of purified S_CoV-2_ proteins in carbonate-bicarbonate buffer (pH 9.6). The next day, plates were blocked with carbonate buffer containing 1% BSA for 2 h at 37°C. Wells were then washed with PBS containing 0.05% Tween 20 (PBS-T), 1:100-diluted sera or 1:10-diluted lung homogenates in PBS-T containing 1% BSA and four serial ten-to-ten dilutions were added and incubated during 2h at 37°C. After PBS-T washings, plates were incubated with 1,000-fold diluted peroxydase-conjugated goat anti-mouse IgG (Jackson ImmunoResearch Europe Ltd, Cambridgeshire, United Kingdom) for 1 h. Plates were revealed by adding 100 µl of 3,3’,5,5’-tetramethylbenzidine chromogenic substrate (Eurobio Scientific). Following a 30 min incubation, reaction was stopped by adding 100 µl of 2N H2SO4 and optical densities were measured at 450nm/620nm on a PR3100 reader.

### NAb Detection

Serial dilutions of heat inactivated sera or clarified lung homogenates were assessed for NAbs via an inhibition assay which uses HEK293T cells transduced to stably express human ACE2 and non-replicative S_CoV-2_ pseudo-typed LV particles which harbor the reporter *luciferase firefly* gene, allowing quantitation of the host cell invasion by mimicking fusion step of native SARS-CoV-2 virus (Sterlin et al., 2020). First, 1.5 × 10^2^ TU of S_CoV-2_ pseudo-typed LV were pre-incubated, during 30 min at room temperature, in U-bottom plates, with serial dilutions of each serum in a final volume of 50µl in DMEM-glutamax, completed with 10% heat-inactivated FCS and 100 U/ml penicillin and 100 µg/ml streptomycin. The samples were then transferred into clear-flat-bottom 96-well-black-plates (Corning), and each well received 2 × 10^4^ hACE2^+^ HEK293-T cells, counted in a NucleoCounter NC.200 system (Chemometec, Denmark) contained in 50 µl. After 2 days incubation at 37°C 5% CO_2_, the transduction efficiency of hACE2^+^ HEK293-T cells by pseudo-typed LV particles was determined by measuring the luciferase activity, using a Luciferase Assay System (Promega) on an EnSpire plate reader (PerkinElmer), as detailed elsewhere (Sterlin et al., 2020). Results are expressed as percentages of inhibition of luciferase activity compared to the maximum of luciferase activity in the absence of NAbs.

Serum samples from COVID-19 cases have been recently described elsewhere (Grzelak et al., 2020). Each participant provided written consent to participate in the study, which was approved by the regional investigational review board (IRB; Comité de Protection des Personnes Ile-de-France VII, Paris, France), according to European guidelines and the Declaration of Helsinki. The study was registered as ClinicalTrials.gov (NCT04325646).

### S_FL_ T-cell epitope mapping

In order to map the immuno-dominant epitopes of S_CoV-2_, peptides spanning the whole S_FL_ (Mimotopes, Australia) were pooled by 16, each containing 15 a.a. residues overlapping by 10 a.a. Peptides were dissolved in DMSO at a concentration of 2 mg/ml and diluted before use at 1 µg/ml for ELISPOT or 2-5 µg/ml for ICS (Intracellular Cytokine Staining) in RPMI−1640 medium supplemented with 10% FCS, 100 U/ml penicillin, 100 µg/ml streptomycin, 1 × 10^−4^ M non-essential amino-acids, 1% vol/vol HEPES, 1 × 10^−3^ M sodium pyruvate and 5 × 10^−5^ M of β-mercapto-ethanol. IFN-γ ELISPOT and ICS assays were performed as described previously (Bourgine et al., 2018; Sayes et al., 2016). For ICS, cells were acquired in an Attune NxT Flow cytometer (Thermo Fisher) and the data were analyzed by FlowJo Software (TreeStar Inc.).

### Generation of Ad5 gene transfer vectors

The Ad5 gene transfer vectors were produced using the ViraPower Adenoviral Promoterless Gateway Expression Kit (Thermo Fisher). The sequence containing CMV promoter, BamH1/Xho1 restriction sites and WPRE was PCR amplified from the pTRIPΔU3CMV plasmid, by use of: (i) forward primer, encoding the attB1 in the 5’ end, and (ii) reverse primer, encoding both the attB2 and SV40 polyA signal sequence in the 5’ end. The attB-PCR product was cloned into the gateway pDONR207 donor vector, via BP Clonase reaction. The *hACE2* was amplified from a plasmid derivative of hACE2-expressing pcDNA3.1, while e*gfp* was amplified from pTRIP-ieCMV-eGFP-WPRE (Ku et al., in revision.). The amplified PCR products were cloned into the pDONR207 plasmid via the BamH1 and Xho1 restriction sites. To obtain the final Ad5 plasmid, the pDONR207 vector, harboring *hACE2* or *gfp* genes, was further inserted into pAd/PL-DEST™ vector via LR Clonase reaction (Figure S7).

The Ad5 virions were generated by transfecting the E3-transcomplementing HEK-293A cell line with pAd CMV-GFP-WPRE-SV40 polyA or pAd CMV-hACE2-WPRE-SV40 polyA plasmid followed by subsequent vector amplification, according to the manufacturer’s protocol (ViraPower Adenoviral Promoterless Gateway Expression Kit, Thermo Fisher). The Ad5 particles were purified using Adeno-X rapid Maxi purification kit and concentrated with the Amicon Ultra-4 10k centrifugal filter unit. Vectors were resuspended and stored à −80°C in PIPES buffer pH 7.5, supplemented with 2.5% glucose. Ad5 were titrated using a qPCR protocol, as described (Gallaher and Berk, 2013).

### Western blot

Expression of hACE2 in the lungs of Ad5::hACE2-transduced mice was assessed by Western Blotting. One million cells from lung cell suspension were resolved on 4 – 12 % NuPAGE Bis-Tris protein gels (Thermo Fisher), then transferred onto a nitrocellulose membrane (Biorad, France). The nitrocellulose membrane was blocked in 5 % non-fat milk in PBS-T for 2 hours at room temperature and probed overnight with goat anti-hACE2 primary Ab at 1 μg/mL (AF933, R&D systems). Following three wash intervals of 10 minutes with PBS-T, the membrane was incubated for 1 hour at room temperature with HRP-conjugated anti-goat secondary Ab and HRP-conjugated anti-β-actin (ab197277, Abcam). The membrane was washed with PBS-T thrice before visualization with enhanced chemiluminescence via the super signal west femto maximum sensitivity substrate (ThermoFisher) on ChemiDoc XRS+ (Biorad, France). PageRuler Plus prestained protein ladder was used as size reference.

### Determination of SARS-CoV-2 viral loads in the lungs

Half of each lung lobes were removed aseptically and frozen at −80°C. Organs were thawed and homogenized for 20 s at 4.0 m/s, using lysing matrix M (MP Biomedical) in 500 µl of ice-cold PBS. The homogenization was performed in an MP Biomedical Fastprep 24 Tissue Homogenizer. The homogenates were centrifuged 10 min at 2000g for further RNA extraction from the supernatants. Particulate viral RNA was extracted from 70 µl of such supernatants using QIAamp Viral RNA Mini Kit (Qiagen) according to the manufacturer’s procedure. Viral load was determined following reverse transcription and real-time quantitative TaqMan® PCR essentially as described (Corman et al., 2020), using SuperScript™ III Platinum One-Step Quantitative RT-PCR System (Invitrogen) and primers and probe (Eurofins) targeting S_CoV-2_ gene as listed in Table S1. In vitro transcribed RNA derived from plasmid “pCI/SARS-CoV envelope” was synthesized using T7 RiboMAX Express Large Scale RNA production system (Promega), then purified by phenol/chloroform extractions and two successive precipitations with isopropanol and ethanol. RNA concentration was determined by optical density measurement, then RNA was diluted to 10^9^ genome equivalents/µL in RNAse-free water containing 100µg/mL tRNA carrier, and stored in single-use aliquots at −80°C. Serial dilutions of this in vitro transcribed RNA were prepared in RNAse-free water containing 10µg/ml tRNA carrier and used to establish a standard curve in each assay. Thermal cycling conditions were: (i) reverse transcription at 55°C for 10 min, (ii) enzyme inactivation at 95°C for 3 min, and (iii) 45 cycles of denaturation/amplification at 95°C for 15 s, 58°C for 30 s. Products were analyzed on an ABI 7500 Fast real-time PCR system (Applied Biosystems). PFU assay was performed as recently described (Case et al., 2020).

### Cytometric analysis of lung cells

Lungs from individual mice were treated with 400 U/ml type IV collagenase and DNase I (Roche) for a 30-minute incubation at 37°C and homogenized by use of GentleMacs (Miltenyi Biotech) mAbs. Cells were filtered through 100 µm-pore filters and centrifuged at 1200 rpm during 8 minutes. Cells were then treated with Red Blood Cell Lysing Buffer (Sigma) and washed twice in PBS. Cells were stained as follows. (i) To detect DC, monocytes, alveolar and interstitial macrophages: Near IR Live/Dead (Invitrogen), FcγII/III receptor blocking anti-CD16/CD32 (BD Biosciences), BV605-anti-CD45 (BD Biosciences), PE-anti-CD11b (eBioscience), PE-Cy7-antiCD11c (eBioscience), BV450-anti-CD64 (BD Biosciences), FITC-anti-CD24 (BD Biosciences), BV711-anti-CD103 (BioLegend), AF700-anti-MHC-II (BioLegend), PerCP-Cy5.5-anti-Ly6C (eBioscience) and APC anti-Ly-6G (Miltenyi) mAbs, (ii) to detect neutrophils or eosinophils: Near IR DL (Invitrogen), FcγII/III receptor blocking anti-CD16/CD32 (BD Biosciences), PerCP-Vio700-anti-CD45 (Miltenyi), APC-anti-CD11b (BD Biosciences), PE-Cy7-anti-CD11c (eBioscience), FITC-anti-CD24 (BD Biosciences), AF700-anti-MHC-II (BioLegend), PE-anti-Ly6G (BioLegend), BV421-anti-Siglec-F (BD Biosciences), (ii) to detect mast cells, basophils, NK: Near IR LD (Invitrogen), BV605-anti-CD45 (BD Biosciences), PE-anti-CD11b (eBioscience), eF450-anti-CD11c (eBioscience), PE-Cy7-anti-CD117 (BD Biosciences), APC-anti-FcεR1 (BioLegend), AF700-anti-NKp46 (BD Biosciences), FITC-anti-CCR3 (BioLegend), without FcγII/III receptor blocking anti-CD16/CD32. Cells were incubated with appropriate mixtures for 25 minutes at 4°C, washed twice in PBS containing 3% FCS and then fixed with Paraformaldehyde 4% by an overnight incubation at 4°C. The cells were acquired in an Attune NxT cytometer system (Invitrogen) and data were analyzed by FlowJo software (Treestar, OR, USA).

### qRT-PCR Detection of inflammatory cytokines and chemokines in the lungs of the mice and hamsters

Lung samples from mice or hamsters were added to lysing matrix D (MP Biomedical) containing 1 mL of TRIzol reagent and homogenized at 30 s at 6.0 m/s twice using MP Biomedical Fastprep 24 Tissue Homogenizer. Total RNA was extracted using TRIzol reagent (ThermoFisher) according to the manufacturer’s procedure. cDNA was synthesized from 4 μg of RNA in the presence of 2.5 μM of oligo(dT) 18 primers, 0.5 mM of deoxyribonucleotides, 2.0 U of RNase Inhibitor and SuperScript IV Reverse Transcriptase (Thermo Fisher) in 20 μl reaction. The real-time PCR was performed on QuantStudio™ 7 Flex Real-Time PCR System (Thermo Fisher). Reactions were performed in triplicates in a final reaction volume of 10 μl containing 5 μl of iQ™ SYBR® Green Supermix (Biorad, France), 4 μl of cDNA diluted 1:15 in DEPC-water and 0.5 μl of each forward and reverse primers at a final concentration of 0.5 μM (Table S2, S3). The following thermal profile was used: a single cycle of polymerase activation for 3 min at 95°C, followed by 40 amplification cycles of 15 sec at 95°C and 30 sec 60°C (annealing-extension step). Mice β-globin or hamster ribosomal protein L18 (RLP18) was used as an endogenous reference control to normalize differences in the amount of input nucleic acid. The average *C*_T_ values were calculated from the technical replicates for relative quantification of target cytokines/chemokines. The differences in the *C*_T_ cytokines/chemokines amplicons and the *C*_T_ of the endogenous reference control, termed Δ*C*_T,_ were calculated to normalize for differences in the quantity of nucleic acid. The Δ*C*_T_ of the experimental condition compared relatively to the PBS-immunized individuals using the comparative ΔΔ*C*_T_ method. The fold change in gene expression was further calculated using 2^−ΔΔ*C*^_T_.

### Lung Histopathology

Samples from the lung were fixed in formalin for at least 7 days and routinely embedded in paraffin. Five µm thick paraffin sections were stained with Hematoxylin Eosin & Saffron (HE&S). Microscopic changes were qualitatively described and when applicable scored semi-quantitatively, using: (i) distribution qualifiers (i.e., focal, multifocal, locally extensive or diffuse), and (ii) a five-scale severity grade, i.e., 1: minimal, 2: mild, 3: moderate, 4: marked and 5: severe.

## Supplemental information titles and legends

**Figure S1. Maps of plasmids used for production of LV encoding S**_**FL**_, **S1-S2 or S1 antigens**.

**Figure S2. Lung histology in mice pretreated with Ad5::hACE2 and inoculated with SARS-CoV-2**. Histological analysis in C57BL/6 mice, pretreated with PBS or Ad5::hACE2, followed by i.n. inoculation of 1 × 10^5^ TCID_50_ of SARS-CoV-2. Analysis was performed at 3 dpi. Lung, HE&S stain, Original magnification: x10, scale bar: 100 µm. Br: Bronchi or bronchiole. Bv: Blood vessel. Arrow: Mononuclear inflammatory cell infiltration.

**Figure S3. Protective potential of systemic immunization with LV::S**_**FL**_ **against SARS-CoV-2 in mice. (A)** Timeline of vaccination by a single i.p. injection of LV followed by Ad5::hACE2 pretreatment and i.n. SARS-CoV-2 challenge. **(B)** Lung viral loads in unvaccinated mice (PBS), LV::S_FL_- or sham-vaccinated mice, at 3 dpi. Statistical significance of the differences in the viral loads was evaluated by two tailed unpaired t test; * = *p*<0.0139.

**Figure S4. Serum or lung homogenate Ab responses against RBD or S1. (A)** Titers of anti-S_CoV-2_ IgG, able to bind to S1 or RBD fragments of S_CoV-2_, as quantitated by ELISA in the sera of C57BL/6 mice primed i.p. at week 0 and boosted i.p. or i.n. at week 3. Titers are determined as mean endpoint dilution. **(B)** Titers of anti-S_CoV-2_ IgG and IgA Abs, as determined in the clarified lung homogenates by ELISA using RBD in coating.

**Figure S5. Inflammation mediators in the lungs of LV::SFL-or sham-vaccinated and challenged mice**. Relative log_2_ fold change in cytokine and chemokine mRNA expression in the challenged “LV::SFL i.p.-i.p.” or “LV::SFL i.p.-i.n.” groups, as determined by qRT-PCR, applied to total lung homogenates prepared at 3 dpi. Relative expressions are normalized to the mean values obtained in PBS-treated unchallenged controls. Shown are inflammatory mediators: **(A)** significantly unchanged between the “LV::SFL i.p.-i.n.” and “sham i.p.-i.n.” groups, **(B)** significantly different between “LV::SFL i.p.-i.p.” and “sham i.p.-i.p.” groups, and **(C)** significantly unchanged between “LV::SFL i.p.-i.p.” and “sham i.p.-i.p.” groups. Statistical significance was evaluated by two tailed unpaired t test; * = p<0.05, ** = p<0.01, *** = p<0.001 and **** = p <0.0001.

**Figure S6. Inflammation mediators in the lungs of LV::S**_**FL**_**-or sham-vaccinated and challenged hamsters**. Relative log_2_ fold changes in cytokines and chemokines mRNA expression in LV::S_FL_-vaccinated and protected hamsters versus unprotected sham-vaccinated individuals, as determined at 4 dpi by qRT-PCR in the total lung homogenates and normalized to the mean of untreated controls. Shown are the inflammatory mediators for which the differences between the LV::S_FL_- and sham-vaccinated hamsters were not statistically different, as evaluated by one-way ANOVA.

**Figure S7. Maps of plasmids used for production of Ad5 encoding hACE2**.

**Table S1. Sequences of primers and probes for SARS-CoV-2 viral load determination**.

**Table S2. Sequences of primers used to quantitate mouse cytokines and chemokines by qRT-PCR**.

**Table S3. Sequences of primers used to quantitate cytokines, chemokines and transcription factor in Syrian golden hamsters.by qRT-PCR**.

