## Supplemental Information for "Intranasal Vaccination with a Lentiviral Vector Strongly Protects against SARS-CoV-2 in Mouse and Golden Hamster Preclinical Models"

### Supplemental Figures

**Figure S1. Maps of plasmids used for production of LV encoding $S_{FL}$, S1-S2 or S1 antigens.**

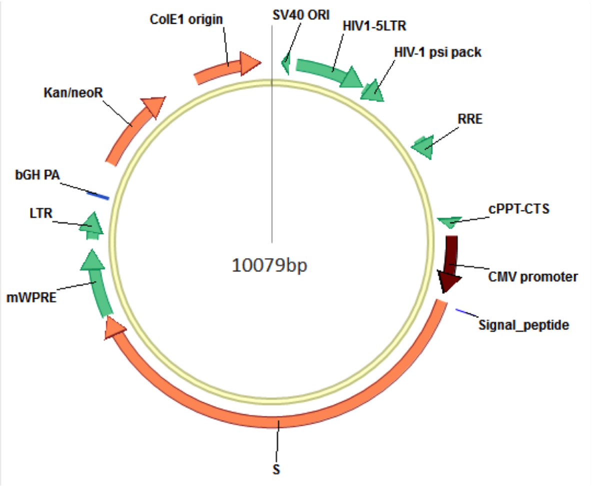


pFlap-CMV-$S_{FL}$-WPREm

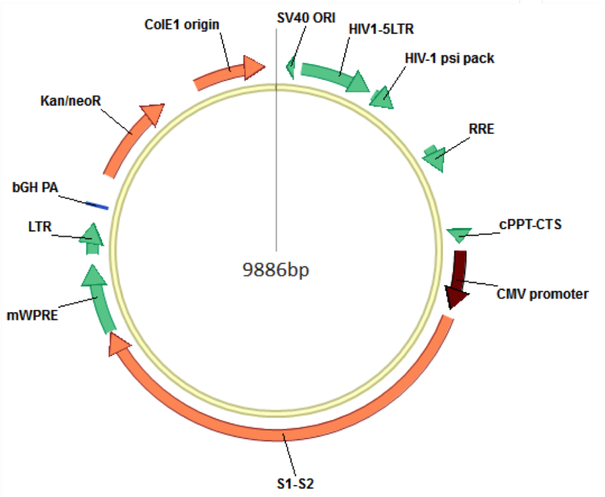


pFlap-CMV-S1-S2-WPREm

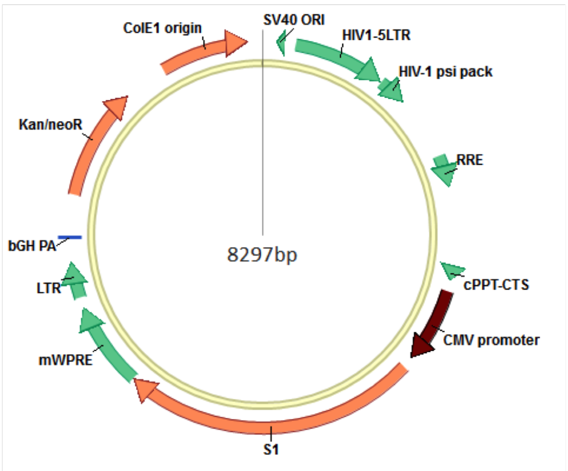


pFlap-CMV-S1-WPREm

**Figure S2. Lung histology in mice pretreated with Ad5::hACE2 and inoculated with SARS-CoV-2.**

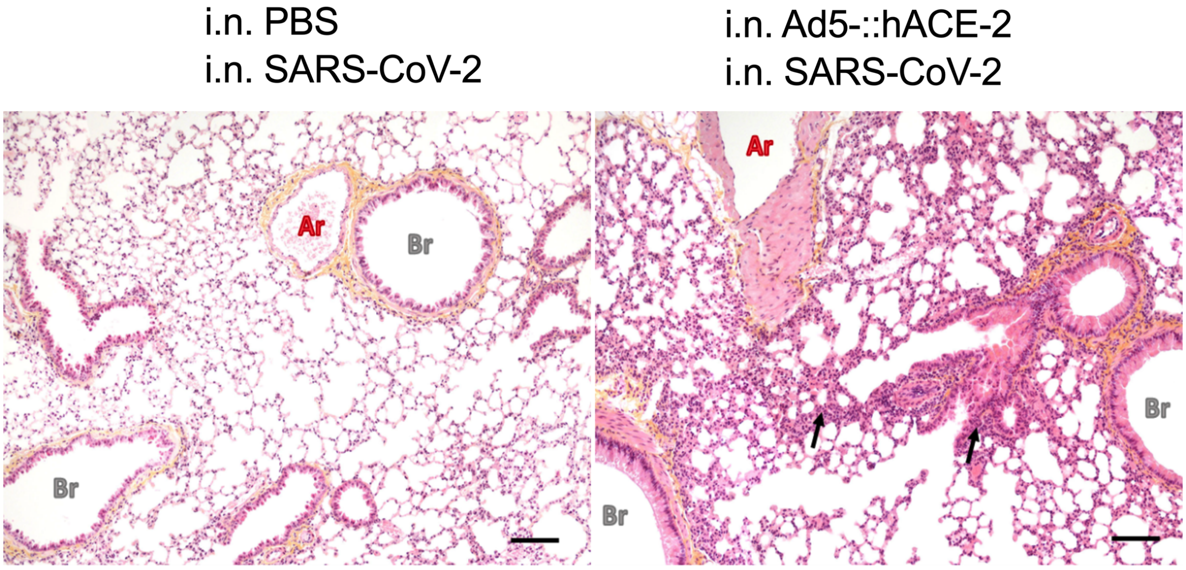

**Figure S3. Protective potential of systemic immunization with LV::$S_{FL}$ against SARS-CoV-2 in mice.**

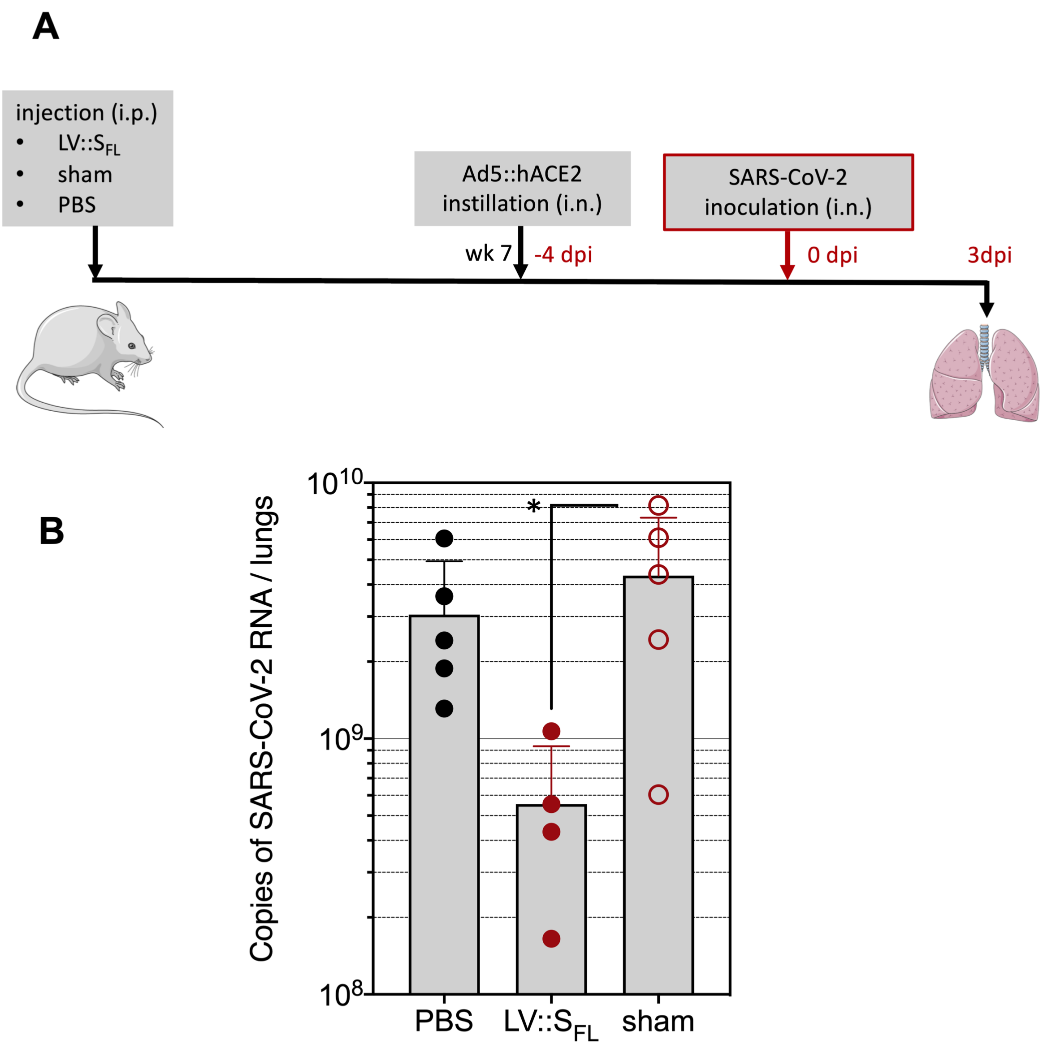

**Figure S4. Serum or lung homogenate Ab responses against RBD or S1.**

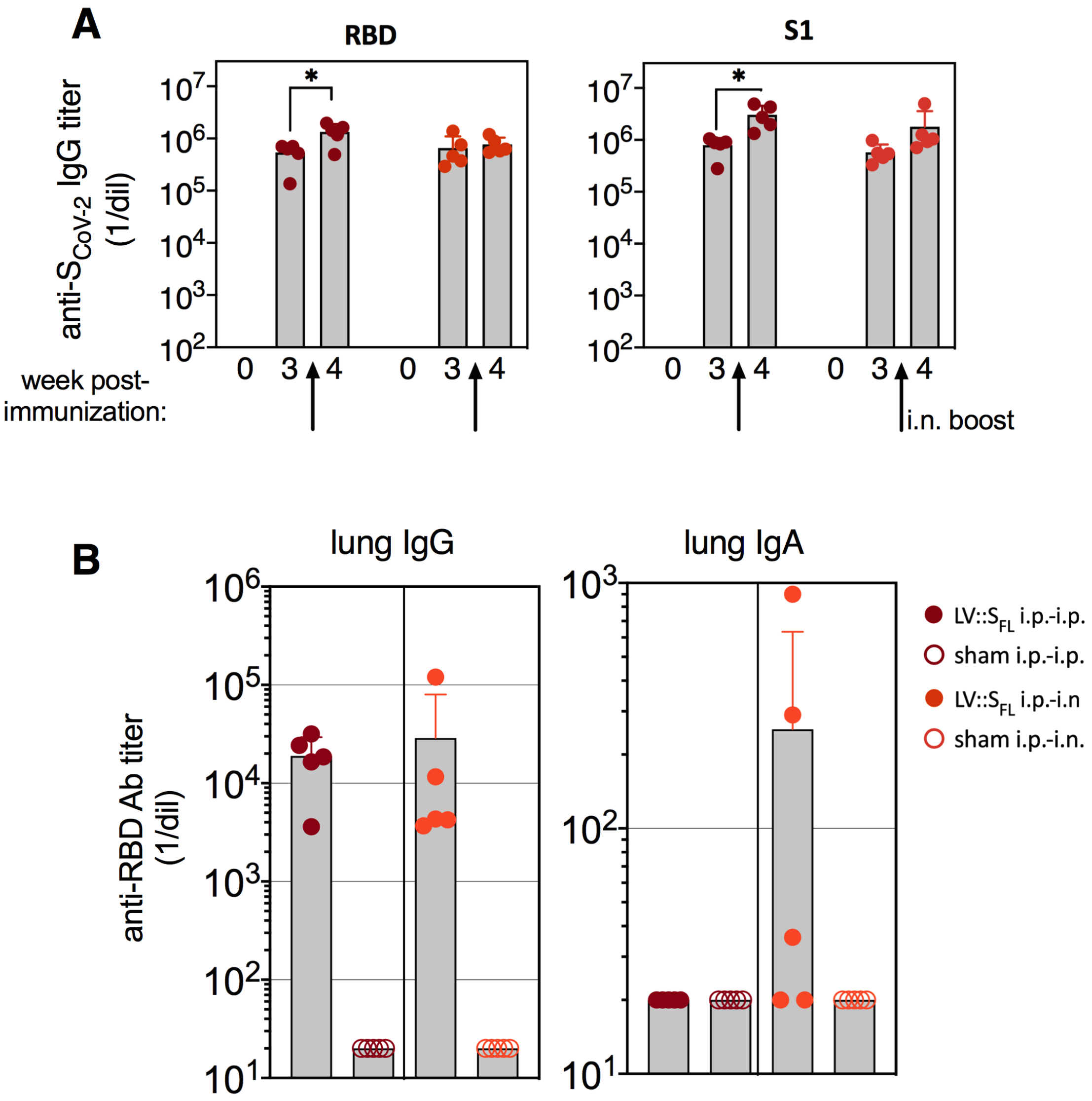

**Figure S5. Inflammation mediators in the lungs of LV::SFL- or sham-vaccinated and challenged mice.**

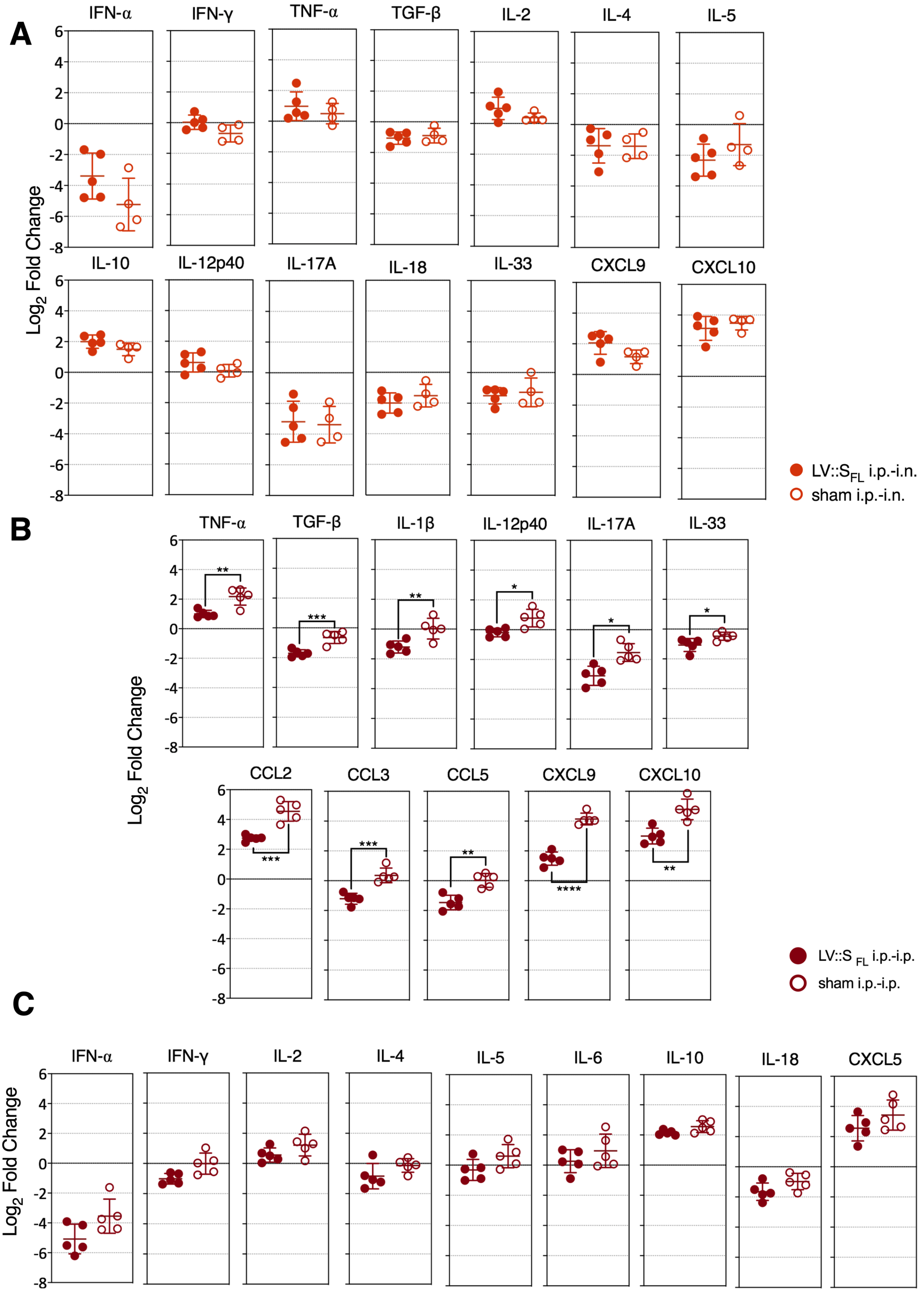

**Figure S6. Inflammation mediators in the lungs of LV::$S_{FL}$- or sham-vaccinated and challenged hamsters.**

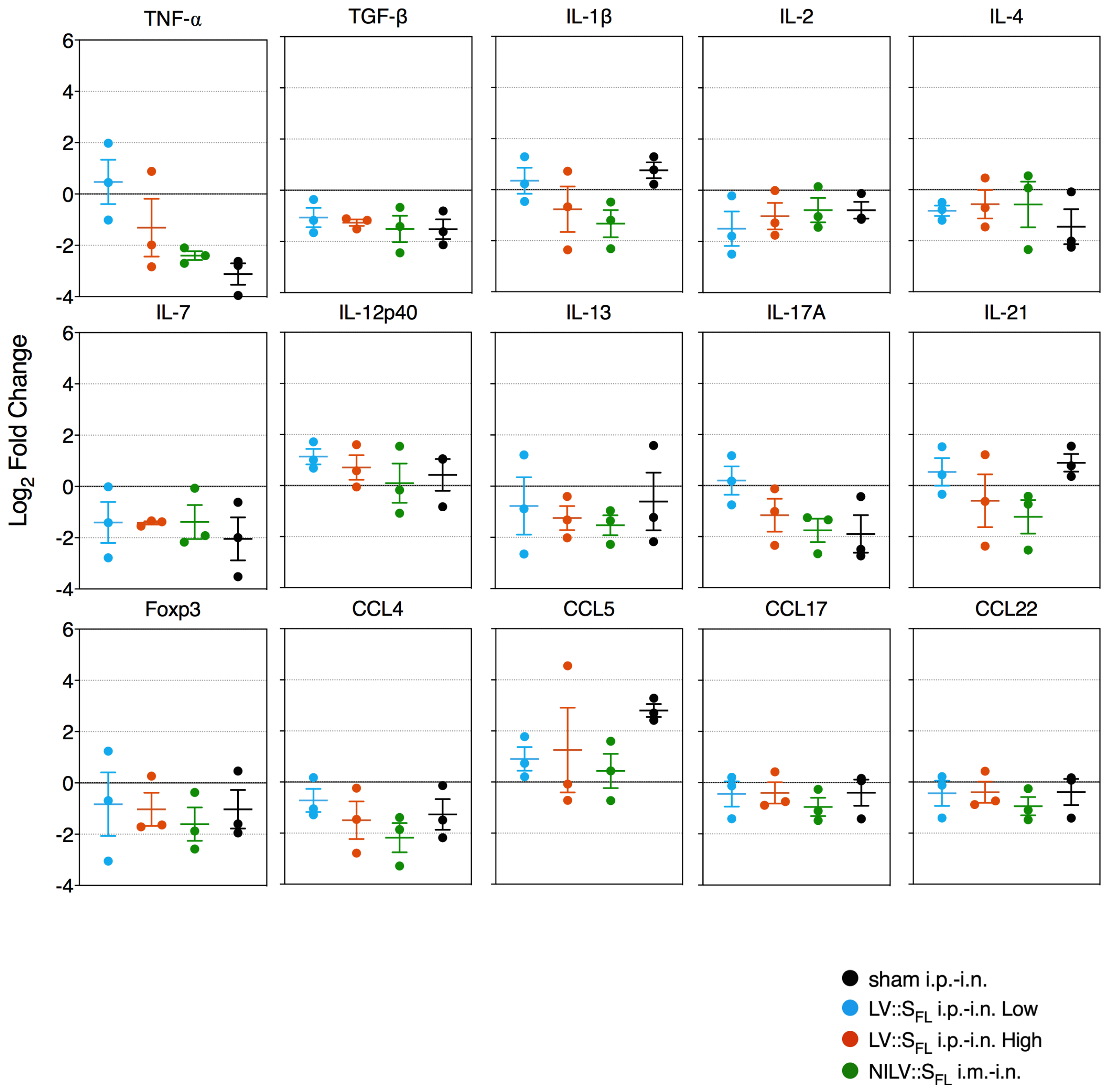

**Figure S7. Maps of plasmids used for production of Ad5 encoding hACE2.**

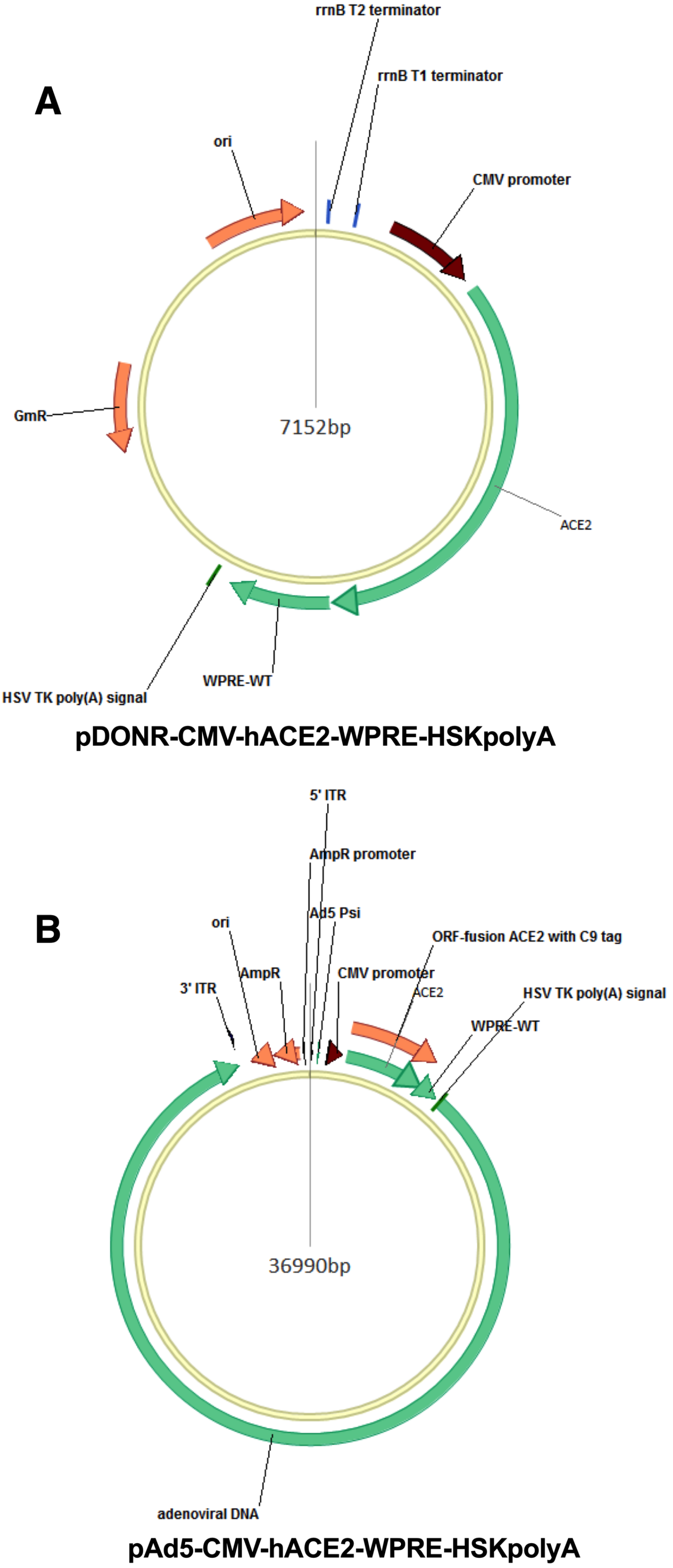

**Table 1S.** Sequences of primers and probes for SARS-CoV-2 viral load determination.

| Primer/Probe Name | DNA Sequences |
|---|---|
| "E-Sarbeco" Fw | 5'-ACAGGTACGTTAATAGTTAATAGCGT-3' |
| "E-Sarbeco" Rv | 5'-ATATTGCAGCAGTACGCACACA-3' |
| "E-Sarbeco" Probe | 5'-FAM-ACACTAGCCATCCTTACTGCGCTTCG-BHQ-1-3' |

**Table S2.** Sequences of primers used to quantitate mouse cytokines and chemokines by qRT-PCR.

| Gene | Sequences |
|---|---|
| β-globin | F : 5'- ATGGGAAGCCGAACATACTG -3' |
| | R : 5'- CAGTCTCAGTGGGGGTGAAT -3' |
| GAPDH | F : 5'- TTCACCACCATGGAGAAGGC -3' |
| | R : 5'- GGCATGGACTGTGGTCATGA -3' |
| IFNα | F : 5'- GGATGTGACCTTCCTCAGACTC -3' |
| | R : 5'- ACCTTCTCCTGCGGGAATCCAA -3' |
| IFNγ | F : 5'- TCAAGTGGCATAGATGTGGAAGAA -3' |
| | R : 5'- TGGCTCTGCAGGATTTTCATG -3' |
| TNFα | F : 5'- CATCTTCTCAAAATTCGAGTGACAA -3' |
| | R : 5'- TGGGAGTAGACAAGGTACAACCC -3' |
| TGFβ | F : 5'- TGACGTCACTGGAGTTGTACGG -3' |
| | R : 5'- GGTTCATGTCATGGATGGTGC -3' |
| IL1β | F : 5'- TGGACCTTCCAGGATGAGGACA -3' |
| | R : 5'- GTTCATCTCGGAGCCTGTAGTG -3' |
| IL2 | F : 5'- CCTGAGCAGGATGGAGAATTACA -3' |
| | R : 5'- TCCAGAACATGCCGCAGAG -3' |
| IL4 | F : 5'- CGAGGTCACAGGAGAAGGGA -3' |
| | R : 5'- AAGCCCTACAGACGAGCTCACT -3' |
| IL5 | F : 5'- GATGAGGCTTCCTGTCCCTACT -3' |
| | R : 5'- TGACAGGTTTTGGAATAGCATTTCC -3' |
| IL6 | F : 5'- CTGCAAGTGCATCATCGTTGTTC -3' |
| | R : 5'- TACCACTTCACAAGTCGGAGGC -3' |
| IL10 | F : 5'- GGTTGCCAAGCCTTATCGGA -3' |
| | R : 5'- ACCTGCTCCACTGCCTTGCT -3' |
| IL12p40 | F : 5'- GGAAGCACGGCAGCAGAATA -3' |
| | R : 5'- AACTTGAGGGAGAAGTAGGAATGG -3' |
| IL17A | F : 5'- GAAGCTCAGTGCCGCCA -3' |
| | R : 5'- TTCATGTGGTGGTCCAGCTTT -3' |
| IL18 | F : 5'- GACAGCCTGTGTTCGAGGATATG -3' |
| | R : 5'- TGTTCTTACAGGAGAGGGTAGAC -3' |
| IL33 | F : 5'- CTACTGCATGAGACTCCGTTCTG -3' |
| | R : 5'- AGAATCCCGTGGATAGGCAGAG -3' |
| CCL2 | F : 5'- AGGTCCCTGTCATGCTTCTG -3' |
| | R : 5'- TCTGGACCCATTCCTTCTTG -3' |
| CCL3 | F : 5'- CCTCTGTCACCTGCTCAACA -3' |
| | R : 5'- GATGAATTGGCGTGGAATCT -3' |
| CCL5 | F : 5'- GTGCCCACGTCAAGGAGTAT -3' |
| | R : 5'- GGGAAGCGTATACAGGGTCA -3' |
| CXCL5 | F : 5'- GCATTTCTGTTGCTGTTCACGCTG -3' |
| | R : 5'- CCTCCTTCTGGTTTTTCAGTTTAGC -3' |
| CXCL9 | F : 5'- AAAATTTCATCACGCCCTTG -3' |
| | R : 5'- TCTCCAGCTTGGTGAGGTCT -3' |
| CXCL10 | F : 5'- GGATGGCTGTCCTAGCTCTG -3' |
| | R : 5'- ATAACCCCTTGG GAAGATGG -3' |

**Table S3.** qPCR primers targeting cytokines, chemokines and transcription factor in Syrian golden hamsters.

| Gene | Sequences |
|---|---|
| β2-Microglobulin | F : 5'- GGCTCACAGGGAGTTTGTAC -3' |
| | R : 5'- TGGGCTCCTTCAGAGTTATG -3' |
| RLP18 | F : 5'- GTTTATGAGTCGCACTAACCG -3' |
| | R : 5'- TGTTCTCTCGGCCAGGAA -3' |
| IFNγ | F : 5'- TGTTGCTCTGCCTCACTCAGG -3' |
| | R : 5'- AAGACGAGGTCCCCTCCATTC -3' |
| TNFα | F : 5'- TGAGCCATCGTGCCAATG -3' |
| | R : 5'- AGCCCGTCTGCTGGTATCAC -3' |
| TGFβ | F : 5'- GGCTACCACGCCAACTTCTG -3' |
| | R : 5'- GAGGGCAAGGAC CTTACTGTACTG -3' |
| IL1β | F : 5'- GGCTGATGCTCCCATTCG -3' |
| | R : 5'- CACGAGGCATTTCTGTTGTTCA -3' |
| IL2 | F : 5'- GTGCACCCACTTCAAGCTCTAA -3' |
| | R : 5'- AAGCTCCTGTAAGTCCAGCAGTAAC -3' |
| IL4 | F : 5'- ACAGAAAAAGGGACACCATGCA -3' |
| | R : 5'- GAAGCCCTGCAGATGAGGTCT -3' |
| IL6 | F : 5'- AGACAAAGCCAGAGTCATT -3' |
| | R : 5'- TCGGTATGCTAAGGCACAG -3' |
| IL7 | F : 5'- ATCAGCATCGATGAATTGGACAAA -3' |
| | R : 5'- CTTGCGAGCAGCACGATTTA -3' |
| IL10 | F : 5'- GGTTGCCAAACCTTATCAGAAATG -3' |
| | R : 5'- TTCACCTGTTCCACAGCCTTG -3' |
| IL12p40 | F : 5'- AATGCGAGGCAG CAAATTACTC -3' |
| | R : 5'- CTGCTCTTGACGTTGAACTTCAAG -3' |
| IL13 | F : 5'- AAATGGCGGGTTCTGTGC -3' |
| | R : 5'- AATATCCTCTGGGTCTTGTAGATGG -3' |
| IL17A | F : 5'- GAGGGAAAGTTGGACCACCA -3' |
| | R : 5'- GACAATGGAGGAAACGCAGG -3' |
| IL21 | F : 5'- GGACAGTGGCCCATA AAACAAG -3' |
| | R : 5'- TTCAACACTGTCTATAAGATGACGAAGTC -3' |
| Foxp3 | F : 5'- GGTCTTCGAGGAGCCAGAAGA -3' |
| | R : 5'- GCCTTGCCCTTCTCATCCA -3' |
| CCL2 | F : 5'- CCGTTAACTCCCCACTCACC -3' |
| | R : 5'- TGAGCTTGGTGATGAAAATCACAG -3' |
| CCL3 | F : 5'- CTCTGAGCCAGGTGTCATTTTC -3' |
| | R : 5'- CCCAGGTTTCTTTGGGGTCA -3' |
| CCL4 | F : 5'- GCTTGGTCACGTGGTCAGTG -3' |
| | R : 5'- GTGGTTGCGCTCCGT GTAG -3' |
| CCL5 | F : 5'- ACTGCCTCGTGTTCACATCA -3' |
| | R : 5'- TTCGGGTGACAAAAACGACT -3' |
| CCL17 | F : 5'- GTGCTGCCTGGAGATCTTCA -3' |
| | R : 5'- TGGCATCCCTGGGACACT -3' |
| CCL22 | F : 5'- TGGTGCCAA CGTGGAAGAC -3' |
| | R : 5'- GAAGAACTCCTTCACTACGCGC -3' |
| CXCL10 | F : 5'- GCCATTCATCCACAGTTGACA -3' |
| | R : 5'- CATGGTGCTGACAGTGGAGTCT -3' |
